# Progressive lifespan modifications in the corpus callosum following a single juvenile concussion in male mice monitored by diffusion MRI

**DOI:** 10.1101/2023.12.21.572925

**Authors:** Andre Obenaus, Brenda P. Noarbe, Jeong Bin Lee, Polina E. Panchenko, Sean D. Noarbe, Yu Chiao Lee, Jerome Badaut

**Author notes:** Corresponding Author: Andre Obenaus, Ph.D. Department of Pediatrics University of California, irvine Hewitt Hall, Rm 2066, Irvine, CA 92697-4475, Office.

## Abstract

**Introduction:** The sensitivity of white matter (WM) in acute and chronic moderate-severe traumatic brain injury (TBI) has been established. In concussion syndromes, particularly in preclinical rodent models, there is lacking a comprehensive longitudinal study spanning the lifespan of the mouse. We previously reported early modifications to WM using clinically relevant neuroimaging and histological measures in a model of juvenile concussion at one month post injury (mpi) who then exhibited cognitive deficits at 12mpi. For the first time, we assess corpus callosum (CC) integrity across the lifespan after a single juvenile concussion utilizing diffusion MRI (dMRI).

**Methods:** C57Bl/6 mice were exposed to sham or two severities of closed-head concussion (Grade 1, G1, speed 2 m/sec, depth 1mm; Grade 2, G2, 3m/sec, 3mm) using an electromagnetic impactor at postnatal day 17. *In vivo* diffusion tensor imaging was conducted at 1, 3, 6, 12 and 18 mpi (21 directions, b=2000 mm^2^/sec) and processed for dMRI parametric maps: fractional anisotropy (FA), axial (AxD), radial (RD) and mean diffusivity (MD). Whole CC and regional CC data were extracted. To identify the biological basis of altered dMRI metrics, astrocyte and microglia in the CC were characterized at 1 and 12 mpi by immunohistochemistry.

**Results:** Whole CC analysis revealed altered FA and RD trajectories following juvenile concussion. Shams exhibited a temporally linear increase in FA with age while G1/G2 mice had plateaued FA values. G2 concussed mice exhibited high variance of dMRI metrics at 12mpi, which was attributed to the heterogeneity of TBI on the anterior CC. Regional analysis of dMRI metrics at the impact site unveiled significant differences between G2 and sham mice. The dMRI findings appear to be driven, in part, by loss of astrocyte process lengths and increased circularity and decreased cell span ratios in microglia.

**Conclusion:** For the first time, we demonstrate progressive perturbations to WM of male mice after a single juvenile concussion across the mouse lifespan. The CC alterations were dependent on concussion severity with elevated sensitivity in the anterior CC that was related to astrocyte and microglial morphology. Our findings suggest that long-term monitoring of children with juvenile concussive episodes using dMRI is warranted, focusing on vulnerable WM tracts.

## Introduction

Concussion awareness has increased considerably over the last decade particularly in sports-related brain injuries (Snyder and Giza 2019). Altered brain connectivity is a signature of concussion injuries both in the acute and in the chronic epochs (Hayes, Bigler et al. 2016, Bouchard, Higgins et al. 2023, Onicas, Deighton et al. 2023). Broadly, these studies report decrements in structural (diffusion magnetic resonance imaging (MRI)) and functional (evoked or resting state MRI) connectivity between affective, motor and somatic brain regions. Pan and colleagues also demonstrated that structural and functional connectivity (coupling) is disrupted in sensorimotor and cognitive circuits after mild traumatic brain injury (mTBI) (Pan, Li et al. 2023). In pediatric mTBI there is sensitivity to white matter tracts, with increased mean diffusivity (MD) in anterior thalamic radiations, arcuate fasciculus and superior longitudinal fasciculus that was dependent on time post injury (Ware, Yeates et al. 2022). The corpus callosum (CC) has significant cortical and intrahemispheric projections and when evaluated in mTBI subjects progressive and considerable alterations were reported in the CC from the subacute to the late chronic (12 month post injury; mpi) time point (Wang, Zhang et al. 2021). Thus, white matter in pediatric and adult mTBI patients is particularly vulnerable to loss of integrity and appears to undergo progressive long-term alterations.

In rodent models, white matter injury has been assessed in several TBI models and at various time points. mTBI induced in mice at different severities reported no behavioral changes within the 14dpi experimental period but increased neurodegeneration in the CC (Velayudhan, Mak et al. 2022). In a mild fluid percussion model (FPI) of brain injury, mice did not exhibit decreased numbers of oligodendrocytes nor their progenitors in the CC at acute 1 or 3dpi time points (Adams, Wood et al. 2023). Interestingly, the authors noted that the number of mature oligodendrocytes were reduced at 3dpi with apparent sparing of myelin albeit suggestions of altered lipid components. CC nodal proteins flanking nodes of Ranvier were also decreased, consistent with presumed altered function (Adams, Wood et al. 2023). Similarly, in a mild repeated cortical contusion injury (CCI) model we reported altered myelin structure at 60dpi with fragmented myelin sheaths (Donovan, Kim et al. 2014). Thus, there are clear alterations to the CC in various models of mild TBI at acute and chronic time points that are likely to impact connectivity.

Clinically, as noted above, non-invasive MRI can track altered white matter composition and connectivity. Diffusion MRI (dMRI) acquired with multiple tensors and with multiple b values can be utilized to assess white matter integrity in rodent models of TBI (as reviewed in (Hutchinson, Schwerin et al. 2018)). A temporal study utilizing the weight drop model in adult mice demonstrated transient dMRI alterations in the CC from approximately 7-21dpi that then resolved by 38dpi (Qin, Li et al. 2018). In adult mice exposed to a mild CCI we reported that by 60dpi axial (AxD) and radial (RD) diffusivity decreased in the CC after a single mTBI, but elevations in these dMRI metrics were observed after repeated mTBI (Donovan, Kim et al. 2014). At even longer time points, Moro and colleagues reported chronic reductions in CC volumes, decreased FA and AxD with increased RD at both 6 and 12mpi in a repeated concussion model (Moro, Lisi et al. 2023). These altered dMRI metrics were coincident with decreased performance in cognition tests at 12mpi, similar to those we previously reported after a single concussion (Rodriguez-Grande, Obenaus et al. 2018). Thus, there is clear evidence morphologically and from dMRI studies that early injury to the CC results in long-term changes in WM microstructure.

However, lacking from the literature are studies that monitor the white matter in the brain longitudinally after a single concussion that mimics the human condition. To address this gap, we developed a model that includes a lateral rotational aspect after a single impact to one hemisphere that was previously well characterized (Rodriguez-Grande, Obenaus et al. 2018). Early MRI changes were noted in the CC up to 1mpi and cognitive alterations were observed at 12mpi (Obenaus, Rodriguez-Grande et al. 2023). To assess the temporal evolution of the dMRI metrics over the lifespan of the mouse, we undertook dMRI at 7 distinct epochs up to 18mpi along with histological assessments of astrocytes and microglia at 1 and 12mpi. Here, we report progressive alterations to the CC after a single concussion to the adolescent mouse (postnatal day 17 injury). These studies provide the basis for future studies investigating therapeutic interventions to ameliorate the advancing white matter decrements in TBI patients.

## Materials and Methods

### Animals

C57BL/6J were bred in–house with breeder mice purchased from Janvier (Le Genest-Saint-Isle, France). Animals were maintained at 21°C ± 1°C, 55% ± 10% humidity, in a 12-h light-dark cycle with access to food and water *ad libitum*. Mild traumatic brain injury (mTBI) was induced on male pups at postnatal day (PND) 17. Males were utilized given the higher incidence of TBI in male children (Coronado, Xu et al. 2011). Animals were randomly assigned (with weight-matching) to a sham group or one of the two mTBI groups with differing severities (Grade 1, G1 or Grade 2, G2) with a minimum sample size of 8 mice per group for a total of n=27-35 per time point. Mice were weaned at PND 25 and housed in groups of 3 to 5 individuals. All animal procedures were carried out following the European Council directives (86/609/EEC), the Animal Research: Reporting of *in vivo* Experiments (ARRIVE) guidelines and local ethics Committee (APAFIS#19296-201902191637994.v4).

### Closed-Head Injury

A mild closed-head injury (CHI) with 2-severity grade (G1, G2) model was utilized as previously described (Rodriguez-Grande, Obenaus et al. 2018, Clément, Lee et al. 2020, Obenaus, Rodriguez-Grande et al. 2023). This clinically relevant model encompasses rotational head movement, requires no surgery to expose the cranium, and incorporates two injury severities to titrate concussive events, better reflecting the inherently heterogeneous nature of mTBI. CHI was administered at PND 17 (Figure 1A). In short, mice were anesthetized using 2.5% isoflurane and 1.5 l/min air for exactly 5 min and then placed on a sheet of tin foil under the impactor tip (3 mm round tip). Without surgery, animals received a focal impact directly over the left somatosensory-parietal cortex center (Bregma approximately −1.7 mm anterior-posterior (AP) coordinate; medial-lateral (ML) coordinate 1.5 mm) using an electromagnetic Leica Impact One Stereotaxic impactor (Leica Biosystems, Richmond, IL, USA) with following parameters: G1= speed=2 m/s, depth=1 mm, dwell time=100 ms; G2 = speed=3 m/s, depth=3 mm, dwell time=100 ms. There were no instances of skull fractures at either severity. Sham animals underwent the same anesthesia procedure and were placed under the impactor apparatus but did not receive an impact. Animals were then allowed to fully recover in individual cages until reappearance of exploratory behaviors and were returned to their original home cage with their respective dam and litter.

**Figure 1.**
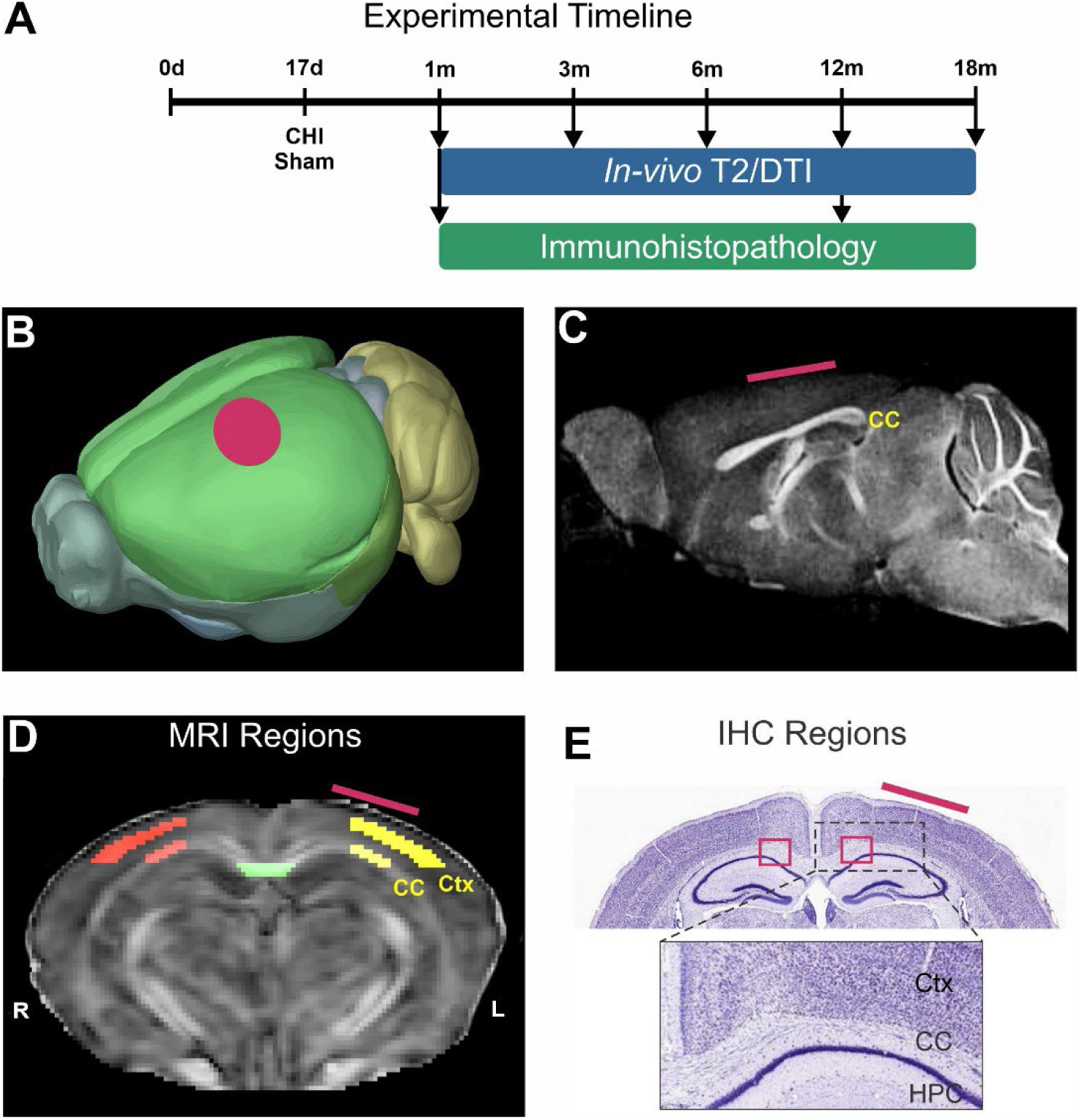
Experimental timeline and regions of interest. A) Experimental timeline with a closed head injury (CHI) at post-natal day 17 illustrates the time points of neuroimaging (from 1 to 18 months post injury, mpi) with correlative immunohistopathology at 1 and 12mpi. B) Schematic brain showing the location of the CHI (red dot). C) Sagittal MRI image demarcating the corpus callosum (CC) location and the location of the CHI (red line). D) MRI regions of interest at the level of the CHI with ipsilateral (yellow) and contralateral (red) cortical (Ctx) and CC regions for analysis. The center CC where the two hemispheres join was also analyzed (green). E) Nissl-stained section illustrating the CC regions situated between Ctx and hippocampus (HPC) analyzed from immunohistochemical sections. (red line indicates location of CHI).

### MR Imaging

Our longitudinal design included 1 (actual age=47d), 3-, 6-, 12-, and 18-months post impact (mpi) time points, involving the same cohort of animals examined *in vivo* throughout all the time points for consistent and comparable outcome measurements (Figure 1A). *In vivo* longitudinal DTI data were acquired on a 7T scanner (Bruker Biospin, Billerica, MA) at each time point with following parameters: repetition time (TR)/ Echo time (TE) = 1000ms/30 ms, 0.267 mm slice thickness, 21 diffusion gradient directions, 1.6 cm × 1.28 cm field of view (FOV), 164×128 acquisition matrix, b=2000 mT/m, and 2 b0 images acquired prior to weighted images. T2-weighted imaging (T2WI) was also acquired with the following parameters: TR/TE=3000ms/7ms, 1.6 cm × 1.28 cm FOV, 162×128 acquisition matrix, 0.8 mm slice thickness, and 25 echoes.

### MR analysis

Acquired MR images underwent preprocessing to optimize image quality and minimize artifacts as previously described (Wendel, Lee et al. 2018). The first echo of T2WI and the mean image of DTI b0s were utilized for brain extraction and registration. The skull was stripped from the brain in a semi-automated manner initially via masks generated using 3D Pulse-Coupled Neural Networks (PCNN3D v1.2) which were then reviewed and adjusted by a blinded experimenter (Chou, Wu et al. 2011). Once the brain was isolated, T2WI data underwent N4 bias field correction with Advanced Normalization Tools (ANTs v2.1). Diffusion data underwent eddy correction and DTI metrics were reconstructed using FMRIB Software Library’s DTIFIT. DTI parametric maps of fractional anisotropy (FA), axial diffusivity (AxD), mean diffusivity (MD), radial diffusivity (RD) and eigenvalues (L1, L2, L3) were obtained. Anisotropy-based shape factor metric components (Weston metrics: linear, planar, and spherical) were derived from FA maps (Benger, Bartsch et al. 2006, Westin, Knutsson et al. 2016, Manninen, Chary et al. 2021).

3D automatic registration was used for brain structure delineation (Clément, Lee et al. 2020). FMRIB’s Linear Image Registration Tool (FLIRT) was utilized to register the Australian Mouse Brain Mapping Consortium (AMBMC) model-based atlas to the T2/DTI native space (Jenkinson, Bannister et al. 2002). The output transformation matrix was then applied to our atlas using ANTs Symmetric Normalization (SyN) algorithm (Jenkinson, Bannister et al. 2002). This robust method not only utilizes many of the same tools and methodologies utilized in clinical MR data processing but allowed for unbiased and rapid segmentation of the corpus callosum and other regions of interest. T2, DTI and volumetric data were extracted using the transformed atlas labels (FSL v5.0; FMRIB, Oxford, UK) (Woolrich, Jbabdi et al. 2009, Andersson and Sotiropoulos 2016).

An additional level of analysis utilized manual regions of interest drawn on the cortex and corpus callosum on 2 contiguous slices centered at the impact site (Figure 1C-D). Delineations were drawn using DSI studio (April 11, 2018 build; http://dsi-studio.labsolver.org). DTI metrics (AxD, MD, RD, FA) were extracted from the regions of interest and summarized in MS Excel.

### Immunohistochemistry (IHC)

At 1 and 12 mpi time points, sham or G2 severity mTBI mice (n=3-4/group) were anesthetized and transcardially perfused with 4% paraformaldehyde (PFA) prepared in phosphate-buffered saline (PBS 1X). Brain tissue was collected and processed as published previously (Clément, Lee et al. 2020, Obenaus, Rodriguez-Grande et al. 2023). Sections of 50 µm thickness were cut using the Vibratome (Leica, Richmond, IL, USA) in the coronal plane and stored at −20°C in a cryoprotective medium (30% ethylene glycol and 20% glycerol in PBS). Two sections per animal (between Bregma −1.3 mm and −2 mm) were used for free-floating IHC. Antigen retrieval was performed to restore the binding of primary antibody to the epitope of interest with sections incubated in a mixture of 1/3 acetic acid and 2/3 of absolute ethanol for 10 min at −20°C and then extensively washed in PBS (6×15 min). To saturate nonspecific antigen-binding sites, the sections were then incubated in blocking solution composed of 1% BSA and 0.3% Triton X-100 in PBS for 2 hours at room temperature (RT). Brain sections were incubated overnight at 4°C with the following primary antibodies diluted in blocking solution: chicken polyclonal anti-mouse GFAP (Abcam ref. ab4674, RRID AB_304558, 1:3000), rabbit polyclonal anti-mouse IBA1 (Wako ref. 019-19741, RRID AB_839504, 1:1500) and rabbit polyclonal anti-mouse NF200 (Sigma-Aldrich ref. N4142, RRID AB_477272, 1:500). The following day slices were washed with PBS (2×10 min) and incubated for 1.5 hours at RT with the corresponding secondary fluorescent antibodies diluted 1:1000 in blocking solution: goat anti-chicken Alexa Fluor^TM^ 568 nm (Invitrogen ref. A-11041, RRID AB_2534098), goat anti-rabbit Alexa Fluor^TM^ 647 nm (Invitrogen ref. A-21244, RRID AB_2535812), goat anti-rabbit Alexa Fluor^TM^ 488 nm (Invitrogen ref. A-11034, RRID AB_2576217). Slices were washed in PBS (3×10 min), mounted on slides and cover-slipped using Vectashield antifade mounting medium with DAPI (Vector laboratories ref. H-1200, Newark, CA, USA) and stored at 4°C.

### IHC Image acquisition

Images were acquired using Nikon eclipse 90i epifluorescence microscope with attached DS-Qi1Mc camera (Nikon Europe, Amstelveen, The Netherlands), pE-300^white^ CoolLED light source (CoolLED, Andover, UK) and NIS-Elements imaging software (Nikon, version 4.30.02). All sections were uniformly stained and clear of background. One coronal brain section per animal was imaged: two z-stacks were acquired with 4X and 20X objectives of the lateral corpus callosum (Figure 1F, Supplementary Figure 1). All acquisition settings of the epifluorescence microscope and software were kept constant within each set of experiments. A negative control staining experiment without the primary antibody showed no detectable signal. Additionally, one z-stack per animal was taken with 40X objective for further fractal analysis.

### Immunohistochemistry analysis

Image analysis was performed using ImageJ software (https://imagej.net/ij/download.html, version 1.54f) (Schindelin, Arganda-Carreras et al. 2012). Immunolabeling intensity was quantified on raw immunofluorescent images taken with 20X objective from ipsilateral and contralateral corpus callosum at the level of the impact site (Supplementary Figure 1). The corpus callosum ROI manually delineated between cingulum bundle (Cg) and hippocampus using neurofilament-200 (NF200) neuronal immunostained sections. The CC ROI was then duplicated from FITC channel (NF200) to Cy3 channel (GFAP, astrocytic marker) and Cy5 channel (IBA1, microglial marker). Mean gray values of the CC ROI was measured at a single focal plane.

### Morphometric analyses

Skeleton analysis was performed on 20x stack acquisitions from GFAP- and IBA1-immunolabeled sections as published (Young and Morrison 2018, Clément, Lee et al. 2020). Briefly, z-stack images were converted from RGB to 8-bit grayscale format and 2D z-projections with maximum intensity were used for analysis. The background noise was removed using Fast Fourier Transform (FFT) bandpass filter (default settings: filter small structures up to 3 pixels, large structures down to 40 pixels, no stripes suppression) and unsharp mask filter (radius of 1 pixel and mask weight 0.9). Resulting images were converted to binary using manual threshold function (black cells on white background). Manual editing was applied to cells in the manually delineated CC ROI to obtain a single set of pixels per cell needed for analysis. Neighboring cells’ structures were removed from analysis using paintbrush tool, cell’s fragmented processes were reconnected where needed by adding missing pixels with free-hand line tool followed by “Fill” function and cells somas were completed using pencil tool. Editing was performed by the same experimenter to avoid inter-user variability. Binary images were then skeletonized where each skeleton was compared to the original cell by creating an overlay on original z-projection to avoid artifacts. Manual correction of skeletons/binary images were performed as required using pencil tool and non-representative skeletons from incomplete cells were removed using paintbrush tool. The final skeletons were analyzed with ImageJ Analyze Skeleton (2D/3D) plugin (Arganda-Carreras, Fernandez-Gonzalez et al. 2010, Clément, Lee et al. 2020) without elimination of endpoints (no prune cycle method). On average, 6-9 IBA1-positive cells (12 mpi time point) and 7-17 GFAP-positive cells (1 and 12 mpi) were analyzed in ipsilateral CC ROI per acquisition. The morphological features analyzed were number of branches and junctions and number of slab voxels (converted from pixels to total process length expressed in µm). For statistical analysis, cells were pooled together within each experimental group (sham or G2).

Fractal analysis was used to assess microglia (IBA1+) in ipsilateral corpus callosum at 12 mpi. We used Fraclac plugin for ImageJ (version 2015Sep090313a9330) and followed previously published protocols (Karperien, Ahammer et al. 2013, Morrison, Young et al. 2017, Young and Morrison 2018, Fernandez-Arjona, Grondona et al. 2019). Z-stacks were acquired with 40X objective for increased process details of microglial cells in ipsilateral (G2) or left CC (sham). Two-dimensional z-projections were filtered and converted to binary. All IBA1-positive cells with clearly visible somata and processes located in CC ROI were analyzed. Manual editing was applied in the same manner as for skeleton analysis. Accurate visual verification was carried out during the editing process to ensure that the shape of the cell on 2D edited binary image corresponded to its original shape on the z-stack.

A batch mode was used in Fraclac plugin (default settings in box counting: binary, no filters, white background locked, 12 grids in grid design, scaling method with default sampling sizes) to analyze all individual cells binary images simultaneously. The convex hull (straight line segments joining the outermost foreground pixels) and bounding circle (the smallest circle around the convex hull) were generated for each cell image by Fraclac plugin. Fractal analysis allows quantification of multiple morphological features of each cell, such as complexity (fractal dimension, D_B_), convex hull span ratio (ratio of major and minor axes of the convex hull; range 0-1), cell area (total number of pixels of the cell converted to µm^2^; one pixel area=0.0276 μm^2^), cell perimeter (expressed in μm; one pixel side=0.1661 µm), cell circularity ((4π*cell area) / (cell perimeter)^2^), density (foreground pixels / total number of pixels in convex hull; range 0-1) and others (Morrison, Young et al. 2017, Fernandez-Arjona, Grondona et al. 2019). Box counting method used by Fraclac determines the amount of pixel detail with increasing scale (Karperien, Ahammer et al. 2013). A higher fractal dimension (D_B_ ranges from 1 to 2) represents a greater complexity of the microglial cell. Effect size was 14 microglial cells pooled from 3 sham mice and 12 microglial cells pooled from 3 G2 mice.

### Statistical analysis

MRI data derived regions over/under the first/third quartile 1.5xIQR (interquartile range) were excluded using Microsoft Excel (Wendel, Lee et al. 2018). GraphPad Prism 7.00 (for Windows, GraphPad Software, San Diego, California USA, www.graphpad.com) was used for data analysis. For comparisons of single features between two groups, a t-test with Welch’s correction was used. For comparisons of a single feature between three groups, one-way ANOVA with Tukey post-hoc test was utilized. For data with multiple time points or with multiple groups and multiple features, a two-way ANOVA with Tukey post-hoc test was used. Pearson’s correlation testing and linear regressions were used to assess correlations between two parameters. The alpha level was set to 0.05. All data are presented as the mean ± standard error of the mean (SEM) unless otherwise indicated.

## Results

We assessed the volume and microstructure of the largest white matter structure in the mouse, the corpus callosum (CC), which was monitored longitudinally across the life span of male mice after a single mild concussive event at postnatal day 17 (PND17) using multimodal magnetic resonance imaging (MRI). Specifically, we undertook diffusion MRI (dMRI) to probe microstructure of the CC, multi echo T2-weighted imaging (T2WI) for regional volumes and T2-relaxation measures to report water content. We also assessed whole cerebrum volumes from MRI. Finally, glial morphological changes were assessed by immunochemistry (IHC) in the CC.

### Volumetric Brain and CC Modifications

Whole brain volumes across the mouse lifespan were not significantly different between shams, Grade 1 (G1) and Grade 2 (G2) mice across time (p=0.652, two-way ANOVA, Age X Group) (Supplementary Table 1). There was a significant effect of time within the G2 group with 1 month post injury (mpi) compared to 3, 6, 12 and 18mpi (p<0.0001, two-way ANOVA). In sham and G1 groups only 1 vs. 18 mpi were significantly different. When ipsi- and contralateral hemisphere volumes were assessed, no significant Age X Group interactions were found; however, significant increases in brain volume at 1mpi in the G1 and G2 groups relative to 18mo. Thus, there were no significant differences in brain volumes between the groups spanning their lifespan.

Hemispheric CC volumes exhibited ipsi- and contralateral differences between groups and summarized in Table 1. The contralateral CC volume showed no significant differences between groups except for G2 mice whose CC volumes were significantly reduced relative to shams at 12mpi. The ipsilateral CC volumes were significantly increased by 6.17% in G1 mice at 3mpi (p<0.05) and the G2 mice at 12mo were reduced by 7.69% at 12mpi (p<0.001) relative to shams. Thus, a mild concussion early in life modulates CC volumes across age.

**Table 1:** Corpus Callosum (CC) volumes (mm^3^) Ipsilateral CC.

| Time point (mpi) | Sham | G1 | G2 | Change G1 | Change G2 |
| --- | --- | --- | --- | --- | --- |
| 1 | 3.87 ± 0.36 | 3.80 ± 0.28 | 3.77 ± 0.22 | -1.80% | -2.65% |
| 3 | 3.86 ± 0.30 | 4.10 ± 0.21 | 3.99 ± 0.17 | 6.17% | 3.29% |
| 6 | 4.19 ± 0.44± | 4.08 ± 0.20 | 4.03 ± 0.23 | -2.68% | -3.82% |
| 12 | 4.21 ± 0.31 | 4.03 ± 0.36 | **3.88 ± 0.09\*** | -4.23% | -7.69% |
| 18 | 4.06 ± 0.37 | 4.13 ± 0.22 | 4.10 ± 0.22 | 1.80% | 0.98% |

\* p<0.05 from Sham

**Contralateral CC**
| Time point (mpi) | Sham | G1 | G2 | Change G1 | Change G2 |
| --- | --- | --- | --- | --- | --- |
| 1 | 3.62 ± 0.14 | 3.75 ± 0.31 | 3.68 ± 0.20 | 3.66% | 1.60% |
| 3 | 3.76 ± 0.27 | **4.08 ± 0.27\*** | 3.97 ± 0.27 | 8.54% | 5.53% |
| 6 | 4.14 ± 0.46 | 4.06 ± 0.25 | 3.93 ± 0.19 | -2.00% | -5.08% |
| 12 | 4.04 ± 0.03 | 4.09 ± 0.35 | **3.82 ± 0.12\*\*\*** | 1.28% | -5.43% |
| 18 | 4.00 ± 0.38 | 4.11 ± 0.26 | 4.01 ± 0.39 | 2.67% | 0.18% |

\* p<0.05 from Sham
\*\*\* p<0.0001 from Sham

### dMRI Microstructure: Fractional Anisotropy (FA)

Two independent analyses were performed, with one examining the average FA across the entire (whole) CC which was then followed by a more focal (region of interest, ROI) examination of the FA in the CC spanning the impact site. In sham mice the contralateral whole CC exhibited increased FA that linearly increased with age starting at 6mpi, in contrast to G1 and G2 mice that exhibited a rapid and significant rise in FA spanning from 1 to 3mpi (*p<0.05) (Fig. 2A). Relative to sham mice the G2 group at 6mpi had a trending increase in FA (p<0.1). The ipsilateral whole CC FA followed a similar pattern but with only G1 mice reporting significant differences relative to shams at 3mpi (#p<0.05) (Fig. 2B). We observed that the G2 group of mice had increased variance in FA in whole CC measures at 18mpi compared to sham and G1 mice (Fig. 2C). This increased heterogeneity of FA within the G2 mice comparative to sham and G1 mice underlies the lack of significant findings at 18mpi despite an average FA decrease of 8.55% in G2 mice compared to shams.

**Figure 2.**
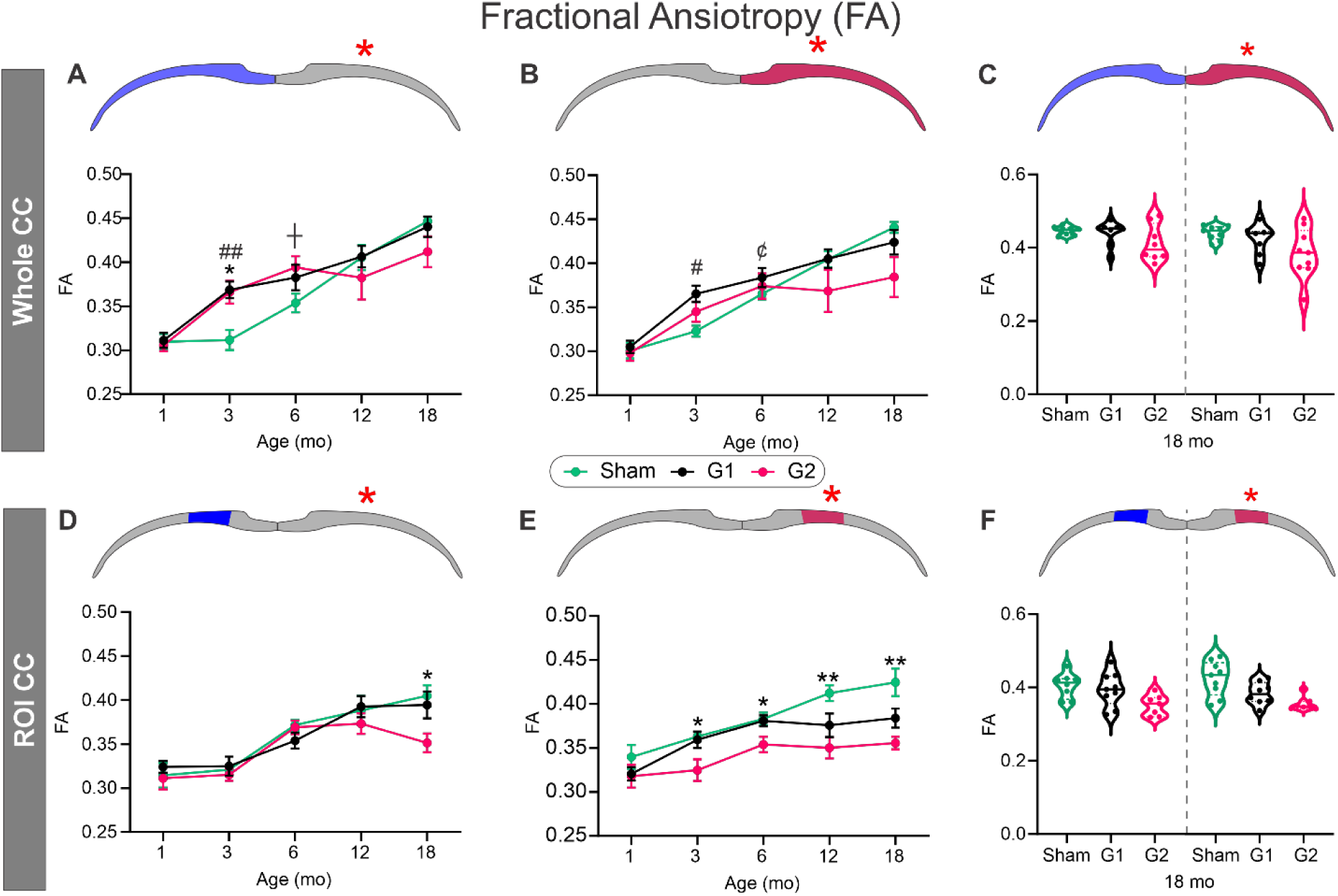
Fractional anisotropy (FA) in the corpus callosum (CC) is reduced late in life after juvenile closed head injury (CHI) at the impact site (red asterisk). A) In the whole CC contralateral to the juvenile CHI both G1 and G2 mice show early elevated FA at 3mpi. In G2, this increase is also trending at 6mpi. By 18mpi G2 FA is reduced relative to the shams. B) The ipsilateral whole CC was elevated at 3mpi in G1 mice, which continues to trend at 6mpi. C) At 18mpi, variance in dramatically increased in the G2, particularly on the ipsilateral injury site. D) In regional measures of the contralateral CC, FA was significantly reduced at 18mpi. E) In contrast, the regional ipsilateral CC FA was significantly reduced at 3-18mpi in the G2 mice, and in G1 at 12 (p=0.101) and 18mpi (p=0.121) after juvenile CHI. F) In the regional CC measures, there was low variance in the FA at 18 mpi compared to whole CC measures (see C). #=p<0.05, ##=p<0.001, ¢=p<0.1 between G1 and S; *=p<0.05, **=p<0.001, ┼p<0.1 between G2 and S.

A different temporal portrait emerged when we undertook regional assessment centered at the impact site. No differences were seen in the contralateral FA with sigmodal temporal evolution (Fig. 2D) in sham, G1 or G2 except for a significant decrease (two-way ANOVA, *p<0.05) in the G2 group at the 18mpi time point. However, a dramatic and significant reduction in FA of the G2 mice was observed spanning the 3-18mpi (two-way ANOVA, 3 and 6mpi, *p<0.05; 12 and 18mpi, **p<0.001) (Fig. 2E). The ipsilateral G1 group also exhibited decrements in FA across the 12 (p-0.100) and 18mpi (p=0.121) time points but did not reach significance. In stark contrast to the variance found in the ipsilateral whole FA at 18mpi, the focal regional FA variance was markedly reduced (Fig. 2F) in the G2 mice compared to sham and G1 mice.

We also examined FA metrics at the midline (center) to determine if this region exhibited increased sensitivity to potential rotational changes in the CC. We observed no overt differences in FA across the 18mpi life span between shams, G1 and G2 mice (Supplementary Fig. 2A). While the G2 group had reduced FA from 6-18mpi, only the final time point was significantly (*p<0.05) reduced relative to sham mice. RD was not significantly different across the 18mpi timeline between shams, G1 and G2 mice (Supplementary Fig. 2B). Post testing revealed a trending increase in RD for G1 at 1mpi (p=0.071) and a significant elevation in G2 at 18mpi relative to shams. Therefore, a single concussion at PND17 elicited evolving decrements in FA particularly within the more severe concussion G2 group. Further, whole CC FA analysis reported global metrics water diffusion asymmetry, while focal CC analyses can yield impactful information about alterations at the injury site, especially in ipsilateral CC.

### dMRI Microstructure: Radial Diffusivity (RD)

RD reports water diffusion perpendicular to the largest diffusion direction and in white matter is thought to reflect axonal and myelin alterations. In whole hemispheric CC analysis in G1 (#) and G2 (*) mice both contralateral (Fig. 3A) and ipsilateral (Fig. 3B) CC RD was exhibited significantly reduced in the ipsilateral G1 and G2 at 3mpi (two-way ANOVA, # p<0.05 and **p<0.001 respectively) and only in G2 at 6mpi (two-way ANOVA, **p<0.001) compared to sham mice. There were no significant differences at 12 and 18mpi despite elevated RD in G2 mice at 18mpi. Virtually identical to FA, at 18mpi there was a large increase in variance in the G2 cohort compared to sham and G1 (Fig. 3C) consistent with broad heterogenous modifications in the CC of G2 mice.

**Figure 3.**
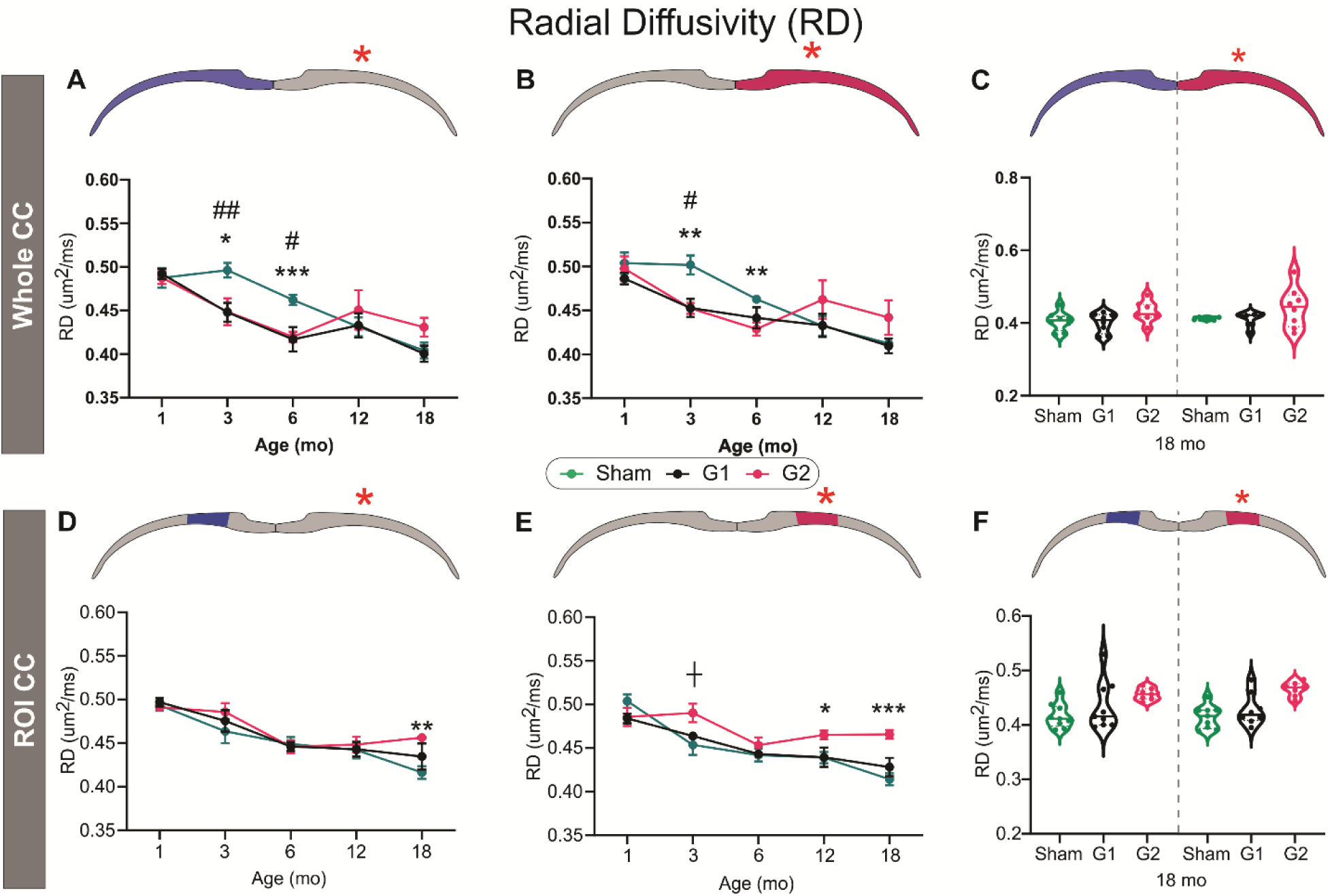
Radial diffusivity (RD) in the corpus callosum (CC) is initially decreased but elevated later in life after closed head injury (CHI) at the impact site (red asterisk). A) The contralateral whole CC RD is decreased at 3 and 6mpi after juvenile CHI in both G1 and G2, but plateaus to sham levels by 12 and 18mpi. B) In the ipsilateral CC a similar trend was observed except the G2 severity elevated at 12 and 18mpi. C) At 18mpi exhibited increased variance in the G2 mice compared to shams in the ipsilateral CC. D) In regional CC measures of the contralateral side, there were no overt differences except at the 18mo time point when G2 RD was elevated relative to Sham mice. E) In the ipsilateral CC a trending increase in RD was observed in G2 mice at 3mpi. At 12 and 18mpi, G2 RD is significantly elevated. F) In regional CC measures the G2 exhibited low variance at 18mpi compared to sham and G1 mice, which also contrasts to the increased variance observed in the whole CC RD measures (see C). # p<0.05, ## p<0.001, between G1 and S; * p<0.05, ** p<0.001, *** p<0.0001, ┼p<0.1 between G2 and Sham.

Focal regional CC analyses demonstrated an entirely different temporal evolution with no changes in RD until 18mpi in the contralateral CC wherein G2 RD is significantly elevated (two-way ANOVA, **p<0.001) (Fig. 2D). Focal ipsilateral CC was significantly increased in G2 at 12mpi (two-way ANOVA, *p<0.05) and at 18mpi (***p<0.001) (Fig. 3E). RD variance in the focal CC analysis at 18mpi was reduced in G2 compared to G1 and sham mice (Fig. 3F). In sum, whole CC analysis showed an early (3-6mpi) elevation in RD while focal CC examination found increased RD at 12-18mpi only in the G2 cohort. This potentially suggests that there are widespread cellular responses to focal concussion but that then leads to more focal altered microstructure within the CC.

### dMRI Microstructure: Mean Diffusivity (MD) and Axial Diffusivity (AxD)

MD represents bulk water diffusion within the CC and is considered a global reporter of microstructural changes. In whole CC assessments significant differences were observed in both ipsi- or contralateral CC at the 3 and 6mpi in both G1 and G2 compared to sham mice(Supplementary Fig. 3A, B). The ipsilateral G1 and G2 mice showed virtually identical reductions in MD at 3 and 6mpi with reductions of ~6% in both ipsi- and contralateral segments compared to shams. No differences were found when regional CC MD was assessed except for G2 mice having increased MD at 18mpi in ipsilateral CC compared to shams (Supplementary Fig. 3C, D).

Axial diffusivity (AxD) has been shown to reflect axonal changes within white matter. Across the 3-18mpi we did not observe any significant changes in AxD in either the ipsi- or contralateral regional CC measures (Supplementary Fig. 4). However, AxD at 1mpi was reduced in G2 mice in both whole or regional CC measures. Thus, AxD results indicate reduced water diffusion at 1mpi while MD is reduced at 3-6mpi (only in whole CC) revealing temporal microstructural sensitivity to dMRI metrics after juvenile concussion.

### Temporal Evolution of Variance and Timeline

We wished to examine the temporal development of the variance we observed in dMRI metrics, particularly FA (see Fig. 2C, F), to gain a deeper insight into how individual mice progressed after concussion with age. Spaghetti plots of FA in sham mice across their lifespan illustrate remarkable homogeneity with only 1 of 10 mice having a lower FA (Fig. 4A). The G1 cohort had modest variance (averaged SEM 1-18mp =0.0102) with most mice clustering around the group mean (Fig. 4B). As shown in Fig. 2C and 3C, the G2 group of mice had robust variance within the CC across the entire ipsilateral CC (averaged SEM 1-18mp =0.0146, a 43.5% increase in SEM) (Fig. 4C). An unexpected finding was the dynamic fluctuations in FA across the lifespan of some G2 mice that likely reflect individual responses to concussion and possibly differential attempts at repair within the CC.

**Figure 4.**
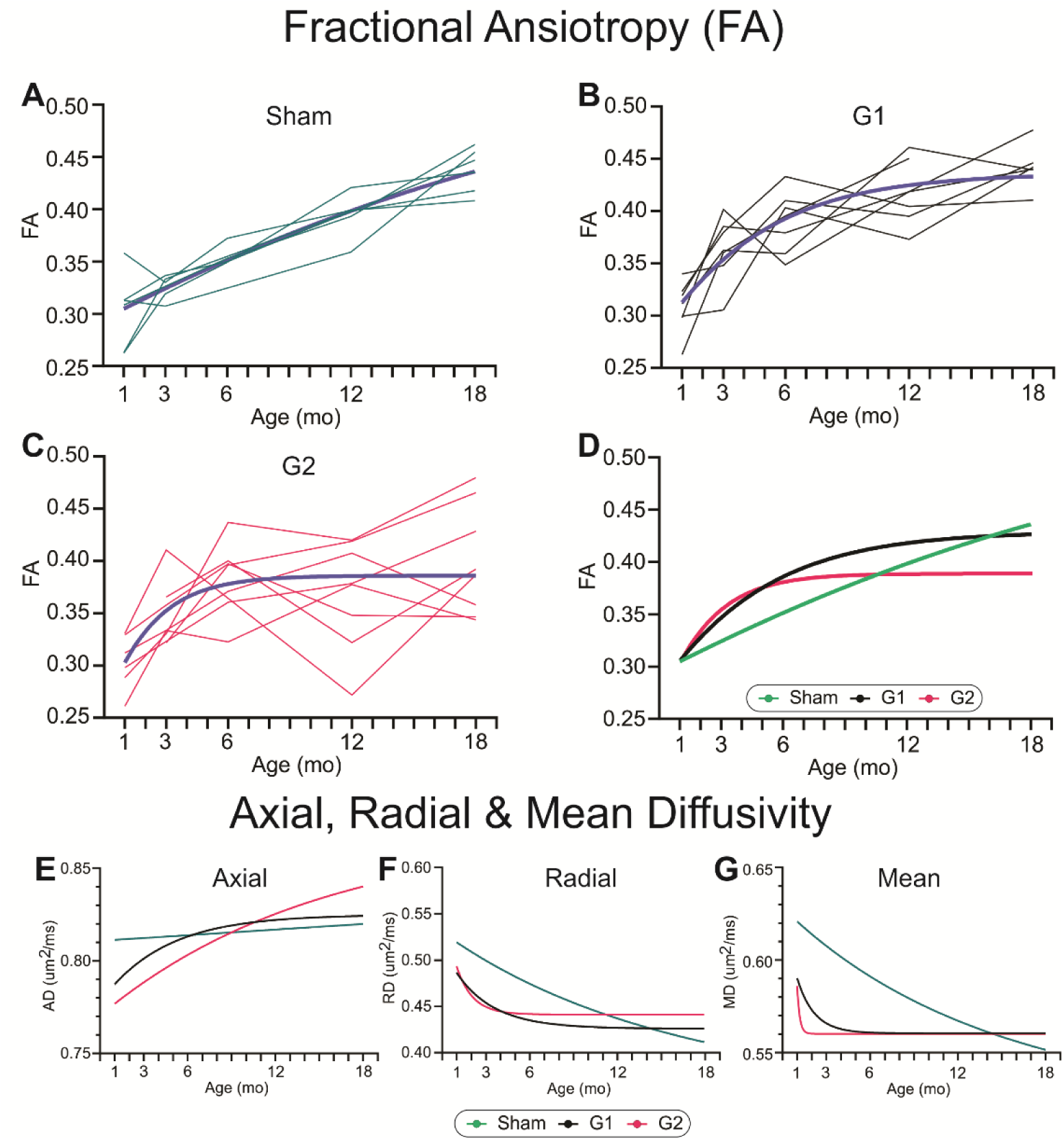
Progression of ipsilateral DTI alterations within the corpus callosum (CC) region of interest (ROI). A) Spaghetti plot of sham animals CC fractional anisotropy (FA) across their lifespan with fitted line, illustrating a linear relationship. B) In G1 mice, there is a curvilinear relationship in the FA of the CC that plateaus by 12mpi. C) Early and sustained FA increases in the CC were observed in the G2 mice that plateaued at 6mpi. D) FA curve fits illustrate the remarkable temporal alterations in the FA of the CC after closed head injury (CHI). E) Axial diffusivity (AxD) was reduced early after CHI in G1 and G2 mice but only G2 mice increased progressively across 18mpi. F) Radial diffusivity (RD) of the CC linearly decreased across the lifespan in sham mice, with both G1 and G2 mice showing rapid decreases early (<3mpi) after CHI that plateaued. G) Mean diffusivity (MD) in shams exhibited a linear decrease but in CHI mice there was a very rapid decline that plateaued and was exacerbated in G2 mice. (A-C – dark blue line represents the fitted curve to the individual mice)

When sham, G1 and G2 cohorts were fitted to a mono-exponential curve (Fig. 4D), the FA in shams showed a linear increase with increasing age. Both G1 and G2 in the months after a single concussive hit exhibited a rapid rise in FA that slowly plateaued to a similar FA level by 18mpi in G1 but was blunted in the G2 mice. A similar analysis was derived for the other dMRI metrics, AxD (Fig. 4E), RD (Fig. 4F) and MD (Fig. 4G). In general, shams showed linear age-related changes across all metrics. AxD was reduced early in G1 but by 3mpi plateaued to sham values which contrasts with the progressive increase in AxD across lifespan in G2. RD in G1 and G2 mice followed a virtually identical trajectory with early decreases that plateaued and were similar to shams at 12mpi. MD timeline was similar to AxD. These analyses suggest that FA and RD appear to be the most sensitive dMRI metrics for monitoring long-term white matter changes.

Whilst the global CC metrics and their variance are informative, we wished to determine where along the CC anterior-posterior extent the variance was derived; in other words, was the variance global or more focal in nature. For these analyses we extracted from each MRI slice the FA along the entire ipsilateral or contralateral CC at 18mpi (Fig. 5) which were then plotted in an anterior-posterior progression as illustrated in Fig. 5A for the ipsilateral CC. In sham mice there was uniform variance across the anterior-posterior extent of the CC in contrast to the robust increased variance in anterior CC at the level of the impact site (red circle on 3D CC reconstruction – center) of G2 mice. The variance was also increased in more posterior aspects of G2 mice as well. We next plotted the average FA in the CC from the contralateral (Fig. 5B) and the ipsilateral (Fig. 5C) hemispheres but found no significant deviations from shams despite the large FA decrease seen at the impact site in G2 mice. Despite the large FA decreases, the lack of significant changes can be attributed to the increased G2 variance.

**Figure 5.**
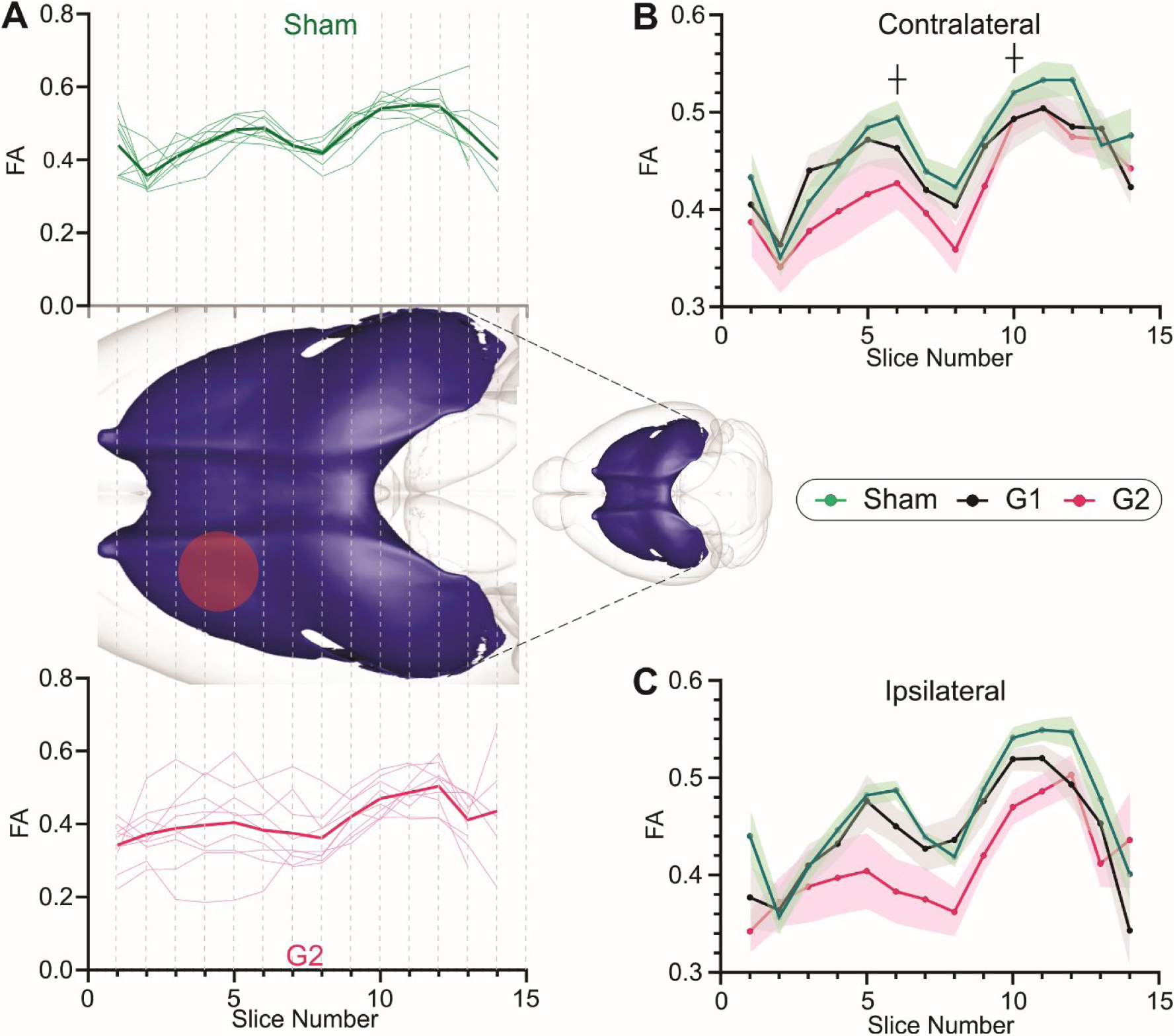
Fractional anisotropy (FA) measures illuminate altered microstructure within the corpus callosum (CC) at 18mpi. A) FA was assessed at each MR slice (slice thickness = 1mm) from anterior to posterior in ipsilateral sham (top panel) and in ipsilateral G2 closed head injury (CHI) mice (bottom panel). FA values were relatively homogenous in the sham mice which starkly contrasts with the large variance in FA at the injury site in G2 mice. Interestingly, posterior to the injury site G2 mice exhibited more homogenous values in FA within the CC. Insert is a 3D reconstruction of the mouse CC where the dotted lines illustrate each MRI slice. The red circle indicates the 3 mm CHI site. B) Contralateral FA values averaged across all cohort mice for sham, G1 and G2 mice illustrate the modest reduction in FA within G2 mice. C) G2 FA values in the ipsilateral CC show a marked decrease in FA compared to G1 and sham mice. ┼ p<0.1 between G2 and Sham.

### Astrocyte and Microglial Morphology after Concussion

From our longitudinal MRI cohort, we extracted 3 G2 CHI and 3 sham mice at selected time points for confirmatory histological study. Our significant findings from the focal CC in FA and RD at 12 and 18mpi prompted us to assess if there were any astrocytic perturbations at the 12mpi time point relative to the 1mpi. Within the contralateral and ipsilateral CC of G2 CHI mice there were no significant differences in glial fibrillary astrocytic protein (GFAP) staining intensity (Fig. 6A) at 1 and 12mpi (Supplementary Fig. 5A) compared to sham mice. We then performed skeleton analysis on astrocytes from the ipsilateral CC to quantify different cell shape features (Fig. 6B). While there were no significant differences in total astrocyte process length between G2 and sham groups at the 1mpi time point (p=0.622), there was a significant decrease (repeated measures t-test, **p<0.01) at 12mpi (Fig. 6C). At 12mpi, sham astrocytes had highly branched skeletal morphology whereas G2 astrocytes had significantly decreased branches and junctions (Supplementary Fig. 5B, C, unpaired t-test, *p<0.05). No significant differences were observed at 1mpi between G2 and sham groups in number of branches (unpaired t-test, p=0.619, Supplementary Fig. 5B) and junctions (p=0.559, Supplementary Fig. 5C).

**Figure 6.**
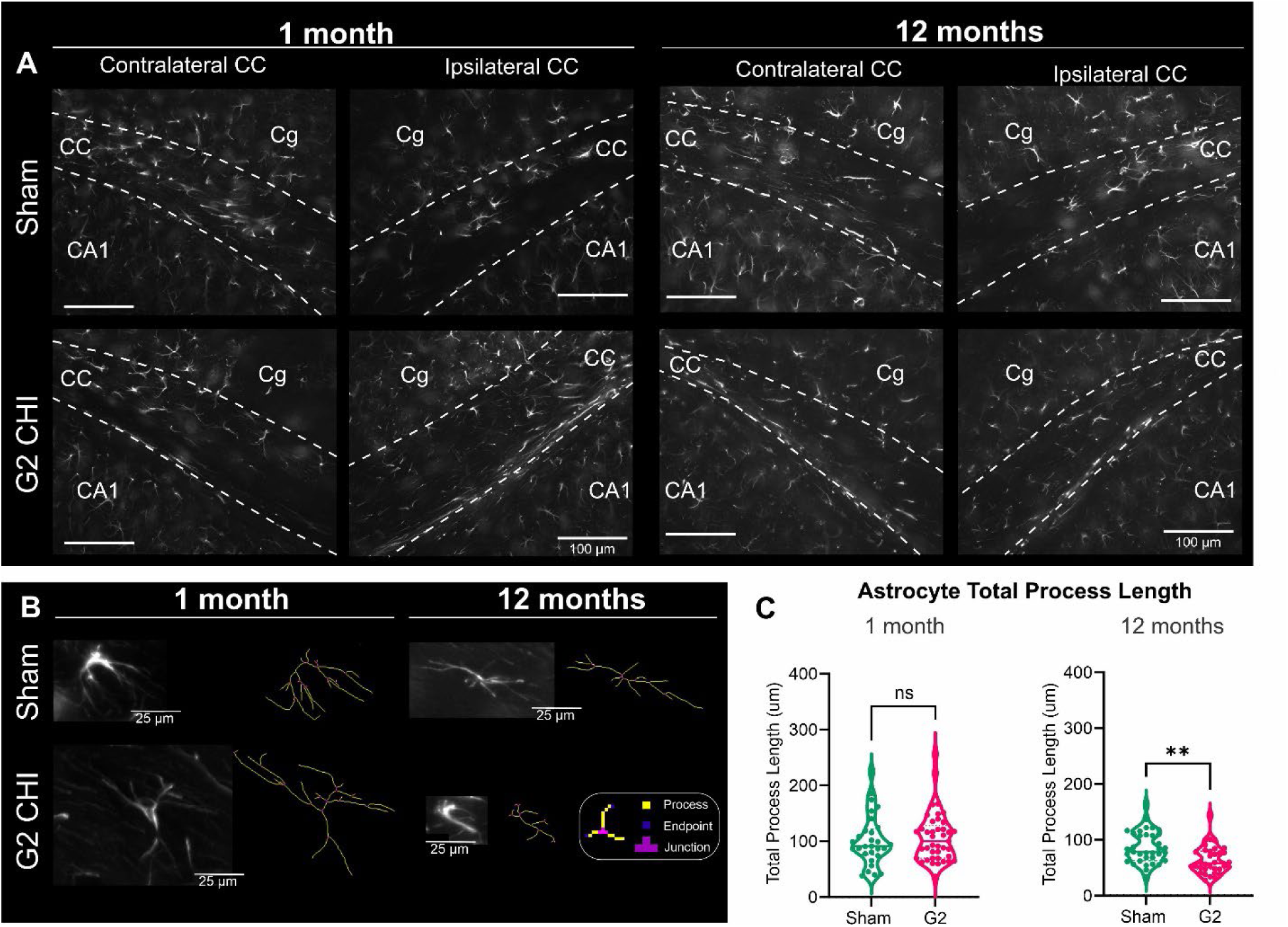
Astrocyte morphology is altered at 12mo after closed head injury (CHI) in the corpus callosum (CC). A) Representative immunohistochemical sections stained for glial fibrillary acidic protein (GFAP) at 1 and 12mpi in shams and in Grade 2 (G2) CHI mice. (Cg – cingulum bundle, CA1 – cornu ammonis 1). B) Sample GFAP-stained astrocytes from the ipsilateral CC illustrate the change in astrocyte morphology with age (1, 12mo) and after CHI. Different morphological cell features are tagged by skeleton analysis as processes (yellow), endpoints (blue) and junctions (magenta). C) Quantitative analyses of the astrocyte total process length (μm) found no significant differences between shams and G2 CHI at 1mo after injury. However, there were significant reductions in astrocyte process length in G2 CHI in the CC at 12mpi compared to shams (** p<0.01, repeated measures t-test). Total process length was measured as the number of slab voxels. See Supplementary Figure 5 for additional astrocyte feature quantification.

Long-term inflammatory responses after TBI have been reported by our group and others (Chen, Islam et al. 2023, Obenaus, Rodriguez-Grande et al. 2023). Given the phenotypic changes in astrocytes we investigated if microglia (via IBA1 immunohistochemistry) within the ipsilateral CC exhibited an activated morphology characteristic of inflammatory states. There were no differences in staining intensity between sham and G2 mice in either the contralateral or ipsilateral CC (Fig. 7A; Supplementary Fig.6A). However, the morphology of G2 microglial cells was altered compared to sham mice with a considerable increase in cellular arborization (Fig. 7B). We quantified these changes using fractal analysis and found that the span ratio of microglial cells was significantly decreased in G2 compared to sham mice in the CC (unpaired t-test, *p<0.05) with a non-significant decrease in cell density which represents the cell size (Fig. 7C, p=0.474). Moreover, the decrease in span ratio coincided with increased cell circularity (unpaired t-test, *p<0.05) suggesting an activated microglial phenotype (Fig. 7C). No significant differences were detected at 12mpi in microglial cell complexity (fractal dimension), cell area and perimeter between sham and G2 group (Supplementary Fig. 6B). Skeleton analysis performed in ipsilateral CC at 12mpi did not show differences in the number of branches or junctions, nor in total process length in microglia (Supplementary Fig. 6C). Taken together, both astrocytes and microglial cells within the CC at 12 months after injury exhibited morphological changes. Neuronal cell staining density was not altered after CHI at either 1 or 12mpi (Supplementary Fig. 7).

**Figure 7.**
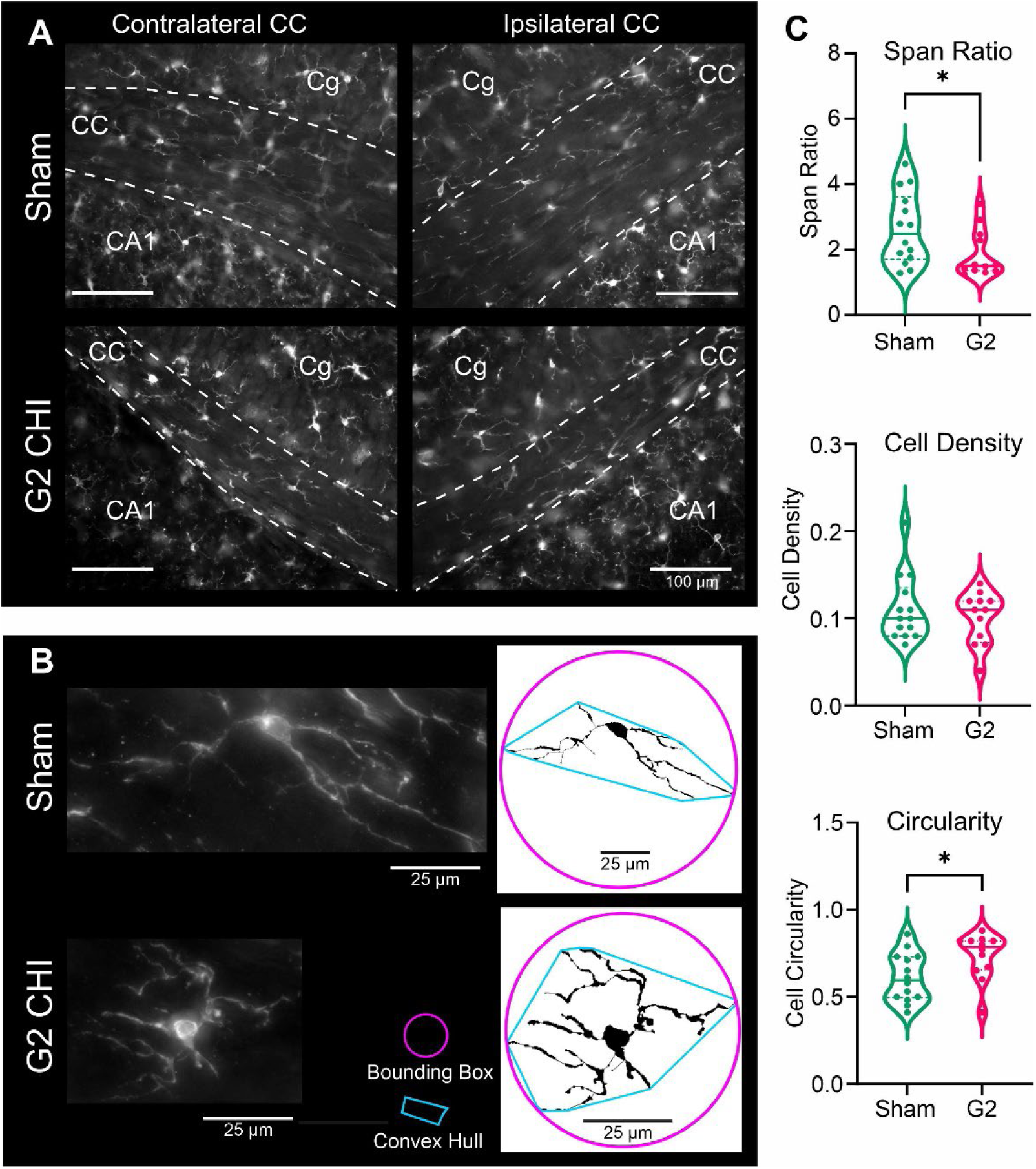
Microglial morphology in the corpus callosum (CC) is altered 12 months after Grade 2 (G2) closed head injury (CHI). A) Representative sections from the contralateral and ipsilateral CC stained with microglial marker ionized calcium-binding adapter molecule 1 (IBA1). (Cg – cingulum bundle, CA1 – cornu Ammonis 1). B) Representative microglia from Sham and G2 CHI mice at 12mpi (left) and its schematic binary image resulting from quantitative fractal analysis (right). Convex hull (blue) and bounding circle (pink) of cells are shown. Note that the Sham microglia schematic is reduced 55%. Cells were pooled from 3 sham and 3 G2 mice. C) Microglial features such as span ratio was significantly reduced with concomitant significant increase in cell circularity of G2 CHI mice (* p<0.05). There were no significant changes in cell density between sham and G2 mice. Thus, CHI induces increased “roundness” of microglial cells in the CC at 12mpi compared to sham mice. See Supplementary Fig. 6 for additional microglial morphometry.

### Fractional Anisotropy (FA) Shape Metrics

Recent studies have reported that FA metrics could be further resolved into linear, planar, and spherical components that provide further insights into the underlying cellular alterations (Westin, Knutsson et al. 2016). Based on our findings of altered inflammatory cell shape and features we undertook FA shape metric analysis. No significant changes in any of the Westin shape metrics were found for contralateral CC (Fig. 8A-C) at 12mpi. In the ipsilateral CC, the linear component was significantly reduced (t-test, *p<0.05) (Fig. 8D) with no change in the planar component (Fig. 8E). The spherical FA component reported a trending increase (t-test, p=0.0513) (Fig. 8F). From these results we hypothesize that water diffusion is more spherical and elongated within the concussed tissues. Thus, in healthy brain tissue (sham) water diffusion is more linear and planar with a reduced spherical shape as it diffuses throughout the CC (Fig. 8G). However, at 12mo after juvenile concussion when astrocytes and microglial morphology is altered, these cellular changes evoke decreased linear (and planar, although not significant) components with increased sphericity (Fig. 8H) as water molecules move through the CC.

**Figure 8.**
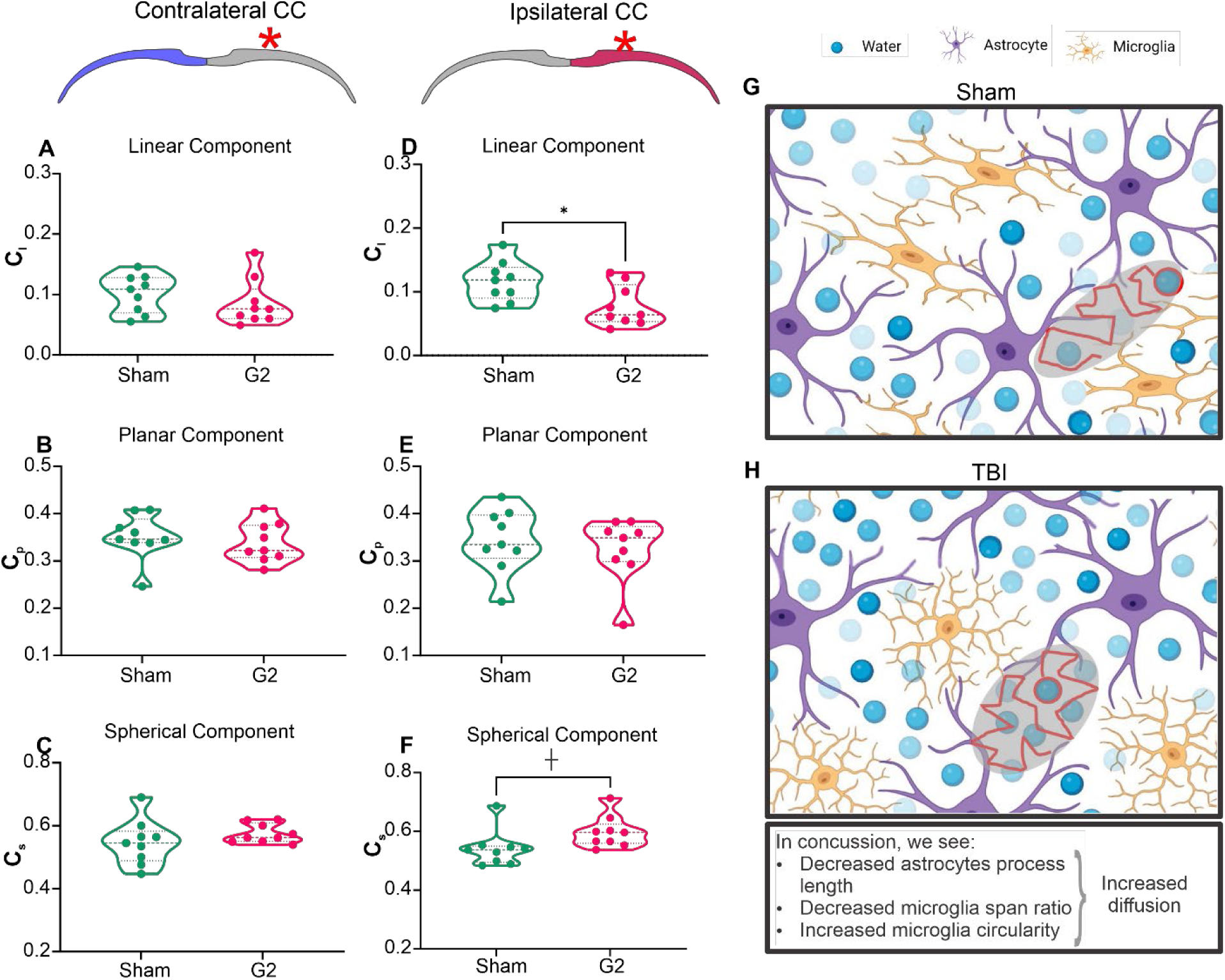
Westin FA shape factor metrics reflect microcellular changes at 12mpi. A, B, C) No significant differences between sham and Grade 2 CHI mice were observed in linear, planar, or spherical components in the contralateral CC. D) The linear component from the ipsilateral hemisphere revealed a significant decrease in G2 mice (* p<0.05). E) No overt changes were observed in the ipsilateral planar component. F) G2 mice showed a trending increase in the spherical component on the ipsilateral side (┼ p=0.0513). G, H) Schematic of how glial changes can influence diffusivity metrics by restricting the movement of water (grey ellipsoids). Specifically, decreased astrocyte process lengths, decreased microglia span ratios and increased microglia circularity suggest increased extracellular space, leading to increased water diffusion between cells, resulting in altered FA shape factor metrics.

## Discussion

Concussion injuries continue to figure prominently in athletic, military, and civilian populations. Studies have alluded to long-term decrements in concussion individuals, including learning and memory, and neuropsychiatric disturbances (de Neeling, Liessens et al. 2023). Critical to understanding the sequala that emerge after concussion are how white matter is modified over time. While progress has been made, long-term pre-clinical studies focused on white matter are lacking. To address this gap, we developed a concussion model of CHI in an adolescent mouse. Sequential dMRI from 1 to 18mpi was performed and analyses focused on the largest white matter tract in the mouse, the corpus callosum (CC). Our novel time course neuroimaging study found: 1) analysis of the whole hemispheric CC in G2 identified decreased AxD at 1mpi followed by elevated FA and decreased RD at 3-6mpi which mirrored the MD decrements, 2) Examination of dMRI metrics at the concussion site reported progressive decrements in FA from 3mpi onwards, with RD being elevated at 12-18mpi only. AxD was decreased at 1mpi followed by decreased RD at 3-6mpi, 3) FA and RD in whole CC across injury groups exhibited increased heterogeneity with increasing severity of concussion, which was predominately localized to the concussion site, 4) Astrocyte and microglial morphology exhibited reactive phenotypes at 12mpi but not 1mpi, and 5) Further analysis of FA using Westin shape metrics found increased spherical and linear components that reflected the inflammatory status of the glial cells. In sum, our findings suggest a progressive temporal vulnerability to the CC after a single juvenile concussion. We also report that a more severe concussion elicits enduring white matter alterations across the lifespan.

The developing brain, in particular white matter, is especially vulnerable to concussion and traumatic brain injury. As reviewed by Semple and colleagues, white matter myelination, synaptogenesis and pruning all are active during adolescence (Semple, Blomgren et al. 2013). Our recently developed mouse concussion model induces injury at post-natal day 17 (PND17) and includes a rotational aspect with early MRI modifications in the CC (Rodriguez-Grande, Obenaus et al. 2018). We also found that two different concussive severities (G1 & G2) resulted in similar perturbations, both on MRI and histopathology, but G2 reported exacerbated CC alterations. On these original studies we next undertook a temporal study spanning 1-18mpi in a cohort of mice that received sham, G1 or G2 concussions using dMRI with glial histopathology assessments. This study is the first to temporally evaluate concussion across the mouse lifespan. A recent adult mouse study found regional CC FA decrements and volumes at both 6 and 12mo but only in repeated mild closed head injury (rmTBI) but not in single TBI mice (Moro, Lisi et al. 2023). In our studies we found significant CC FA changes in G2 mice as early as 3mpi and in G1 mice at 12 and 18mpi when regional analyses were performed. When whole CC analysis was performed only the 3 and 6mpi had group-wise differences, due in part to the heterogeneity of FA changes across the entire hemispheric CC. The differences between Moro et al and our findings can be attributed to concussion model differences; our model explicitly has a rotational component (Rodriguez-Grande, Obenaus et al. 2018).

Several studies have reported on the neuroimaging alterations within the CC at early chronic epochs. Parent and colleagues noted reduced CC FA at 2mpi in rats after a diffuse fluid percussion injury (FPI) during late adolescence (PND31) (Parent, Li et al. 2019). In an open skull cortical contusion model (CCI) mild TBI model we found reduced MD (but not FA) at 2mpi which correlated to increased myelin components in anterior CC (Wendel, Lee et al. 2018). In a young adult rat rmTBI model there was reduced FA in the CC at 2.5mpi with no attendant changes in AxD, RD or MD (Yu, Shukla et al. 2017). Another study in a moderate single TBI (CCI) at 1mpi there were widespread FA reductions in the CC that were evident as early as 1wk after injury, in part due to the more severe nature of the TBI model (Harris, Verley et al. 2016). Thus, despite differences in models utilized, regions of interest, and methods of analyses, there is strong evidence for early and sustained vulnerability to concussive injury, particularly to the corpus callosum. Indeed, many studies have noted cognitive deficiencies in rodent concussion models even early after injury suggesting that a linkage between behavior and white matter integrity after concussion (Yu, Shukla et al. 2017, Moro, Lisi et al. 2023, Obenaus, Rodriguez-Grande et al. 2023).

A novel aspect of the present study is that we were able undertake a longitudinal dMRI study within same mice across 18mpi. These longitudinal studies are difficult, expensive and time consuming, but they provide an exceptional perspective of how white matter in each subject (mouse) progresses across their lifespan. One facet we observed in our temporal studies was the increased variance in dMRI metrics, particularly in the G2 group (Fig. 4). Sham and G1 mice across their lifespan had a relatively consistent variance whilst the G2 group had considerable inter-mouse variability in the FA across their lifespan. Indeed, considerable variance has also been noted in adult and pediatric clinical populations (Oehr, Yang et al. 2021, Stillo, Danielli et al. 2021, Palacios, Yuh et al. 2022, Ware, Yeates et al. 2022). One suggestion to limit this variability is to use symmetry measures to blunt TBI-associated effects on FA (Vakhtin, Zhang et al. 2020). Alternatively, this variance has been suggested to represent individual resilience and may relate to recovery (or lack) after TBI when examining white matter tracts (Schmidt, Lindsey et al. 2021, Cai, Brett et al. 2022). Future work in animal models of concussion should pursue the linkage between variance in dMRI metrics and behavioral outcomes. We have previously reported associations between FA in the hippocampus and cognitive tests (Obenaus, Rodriguez-Grande et al. 2023).

Not all dMRI metrics are created equal nor report the same microstructural details longitudinally. In our study we observed that FA was the most robust reporter of CC alterations over time, particularly in the focal regional CC analyses (Fig. 2). At earlier time points AxD, RD and MD selectively model changes in the white matter microstructure, whereas FA increasingly in the regional CC metrics identifies abnormalities. Several reviews encompassing dMRI and TBI have also summarized that FA is indeed a sensitive potential biomarker (Delouche, Attye et al. 2016, Turner, Lazarus et al. 2021), although there are conflicting reports as well. Monitoring FA, for example at the injury site, could provide the basis for clinical diagnostics related to progression and recovery as well as assessing future therapeutics. Clinically, FA (as well as other metrics) have been shown significant sensitivity to mTBI in patients (Palacios, Yuh et al. 2022). Our findings and those from the literature advocate for monitoring FA closely in mTBI and concussion patients.

The physiological, cellular and molecular alterations that underlie the progressive changes in the CC after a concussion remain to be fully elucidated. A primary focus has been the inflammatory cascade that ensues after concussion particularly in the acute and sub-acute epochs (Verboon, Patel et al. 2021). One year after rmTBI in the CC histopathological evidence of axonal injury is evident, with a 51% reduction in myelin, increased astrocytes (116%), and microglia (69%) (Moro, Lisi et al. 2023). In our study an interesting finding was that the anterior CC was more affected than posterior regions. This is perhaps not overtly surprising as these alterations were found at the concussion site and possibly reflects head rotation in our concussion model. We previously reported that at 1, 7 and 30d post injury there was increased glial fibrillary acidic protein (GFAP; astrocytes) staining in this concussion model within the CC (Rodriguez-Grande, Obenaus et al. 2018). In our current study we did not observe any overt astroglia changes at 1mpi and by 12mpi in CC, but there was a modest but significant reduction in astrocyte branching and process lengths suggesting a reduced spatial coverage. Another study showed a process shortening in CC fibrous astrocytes in response to axonal injury (stab wound) followed by process re-extension and glial scar formation (Sun, Lye-Barthel et al. 2010). Functional role of these morphological changes in astrocytes in response to brain injury and whether these changes are linked to a specific inflammatory state remain unknown. The modulation of myelin dysfunction/repair by astrocytes has not been studied in TBI but research in multiple sclerosis models have observed a dynamic communication between glia and suggests an avenue of future research (Schroder, Mulenge et al. 2023).

Early and acute microglial responses in TBI and concussion have been well documented in our studies and by others (Shitaka, Tran et al. 2011, Taib, Leconte et al. 2017, Jullienne, Hamer et al. 2020, Mohamed, Corrigan et al. 2020). IBA1 immunolabeling in our current study showed a more ameboid microglial cell morphology with reduced span ratio and increased circularity in the more severe G2 concussion mice. We did not find increased microglial density in our concussion study at 12mpi, but others have reported increased CC microglial density at the same time point which is likely due to repeated TBI model (Moro, Lisi et al. 2023). In a study of human TBI there was evidence of neuroinflammation up to 17yrs post injury (Johnson, Stewart et al. 2013). Microglia that contain lipofuscin exhibited increased phagocytic activity in aged compared to young mice and TBI in old mice (18mo) elicited a robust phagocytic response in a sub-population of microglia despite no differences in the number of microglia (Ritzel, Li et al. 2023). Despite the paucity of studies evaluating the long-term microglial response after concussion and TBI, there appears to be ongoing microglial inflammation years after injury, as highlighted in this recent review (Wangler and Godbout 2023). Modulation of oligodendrocytes by microglia can impact myelination after TBI as a potential mechanism to improve outcomes (Song, Hasan et al. 2022).

An intriguing and novel finding of our current study is that further dissection of FA metrics could point to underlying cellularity after concussion. In our G2 concussed mice at 12mo we assessed Westin shape metrics from FA to identify linear, planar and spherical components (Benger, Bartsch et al. 2006). Our analysis of the FA found increased sphericity in the CC that coincides with the increased circularity in microglia and blunted branching patterns in astrocytes. The reduced FA linearity in the CC also tangentially confirms the increased ameboid shapes of both astrocytes and microglia. While additional confirmation needs to be undertaken, these FA shape analyses may in future provide a non-invasive metric to assess the inflammatory status of white matter. Diffusion MRI can be combined with positron-emission tomography (PET) for microglial markers (Masdeu, Pascual et al. 2022). An example of combinatory imaging in severe TBI was reported by Missault and colleagues (Missault, Anckaerts et al. 2019), where they showed increased inflammation (PET) and acute reductions in FA (MRI) that were correlated with behavioral decrements. Future combinatory studies should also evaluate fluid biomarkers for relationships between FA, PET and inflammatory biomarkers.

There are several limitations in the current study. This longitudinal cohort was limited to males and ideally should have included female mice as well. Moro found no sex differences in their long-term rmTBI studies (Moro, Lisi et al. 2023). While sex differences have been reported in TBI (see review (Biegon 2021)), no overt consensus has yet emerged, particularly with regard to adolescent concussions. The current study would be strengthened with a larger longitudinal cohort that would allow cross-sectional extraction of sufficient mice at each time point for immunohistochemical and molecular quantifications. In addition, new emerging advanced dMRI approaches such as multi-shell dMRI, can provide fresh non-invasive insights into the pathophysiology of white matter (McCunn, Xu et al. 2021, Lima Santos, Kontos et al. 2022).

## Conclusions

The clinical and neuropsychological consequences of adolescent concussion are starting to emerge. However, there is a gap in the preclinical literature with a lack of long-term assessments using clinical metrics, such as dMRI, to probe white matter vulnerability after adolescent brain injury. In our study, we provide the first long-term evidence of altered white matter microstructure, where early (1mpi) AxD reminiscent of axonal damage followed by myelin decrements as reported by RD. FA was altered as early as 3mpi and continued to consistently be reduced across the lifespan of the concussed mouse to 18mpi. Interestingly, FA showed a decreased linear component in concussed mice at 12mpi. Brain injury in adolescent mice had a long-term impact on astrocytes which had less complex morphology. Microglial cellular morphology in white matter was consistent with an increased phagocytic/inflammatory phenotype. Another key finding was that the more severe concussion modality exhibited increased variability within the CC and may represent individual mice that might be attempting endogenous recovery of white matter. Future work should utilize this information to guide therapeutic interventions. In summary, we demonstrate progressive and enduring changes within the mouse CC across its lifespan following a single concussion overlying the somatosensory cortex. This study also highlights the paucity of longitudinal studies in both sexes and additional efforts should be undertaken to fill this gap in knowledge.

## Declarations

### Ethics approval

All animal procedures were carried out in accordance with the European Council directives (86/609/EEC) and the ARRIVE guidelines.

### Data Availability

All protocols, raw, and summary data are available upon reasonable request to the corresponding author.

## Acknowledgements

The authors would like to acknowledge

