## Supplemental Table and Figures for "Progressive lifespan modifications in the corpus callosum following a single juvenile concussion in male mice monitored by diffusion MRI"

### Supplementary Figures and Tables

#### Supplementary Table 1A: Number of mice at each time point

| Time point (mpi) | Sham | G1 | G2 |
|---|---|---|---|
| 1 | 12 | 13 | 10 |
| 3 | 9 | 11 | 10 |
| 6 | 10 | 11 | 11 |
| 12 | 9 | 11 | 10 |
| 18 | 9 | 8 | 10 |

mpi = months post injury

#### Supplementary Table 1B: Cerebrum volumes across lifespan

| Time point (mpi) | Sham | G1 | G2 |
|---|---|---|---|
| 1 | 262.08 ± 6.85 | 311.50 ± 9.42 | 262.77 ± 7.55 |
| 3 | 287.24 ± 6.61 | 303.33 ± 4.13 | 297.60 ± 4.55 |
| 6 | 330.30 ± 4.71 | 319.26 ± 3.27 | 321.09 ± 2.60 |
| 12 | 324.47 ± 3.36 | 324.37 ± 2.45 | 317.19 ± 4.19 |
| 18 | 337.28 ± 4.07 | 333.62 ± 3.13 | 329.54 ± 3.91 |

mpi = months post injury

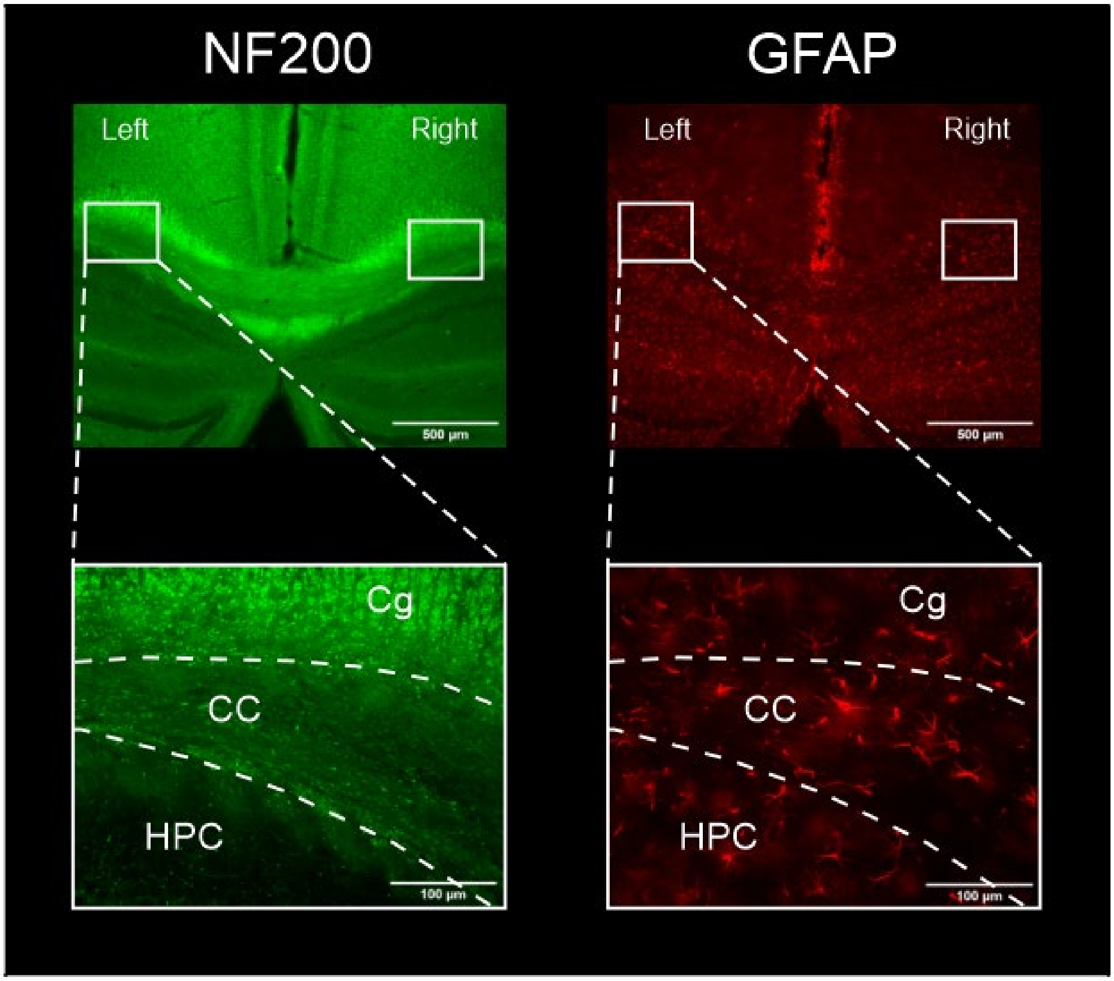


**Supplementary Figure 1. Regions of interest (ROI) in lateral corpus callosum (CC) used for quantitative analyses.** Images of corpus callosum were acquired from immunofluorescent histological coronal sections with 4X (top) and 20X objectives (bottom) in both hemispheres for each animal. Sham mouse at 12mpi time point is represented on this figure. Approximate imaged regions are indicated by white rectangles. Corpus callosum (CC) ROI used for quantitative analyses was manually delineated first on NF200 images (left) and then copied on GFAP image (right). Scale bars: 500 µm for 4X magnification and 100 µm for 20X magnification. Cg – cingulum bundle, HPC – hippocampus.

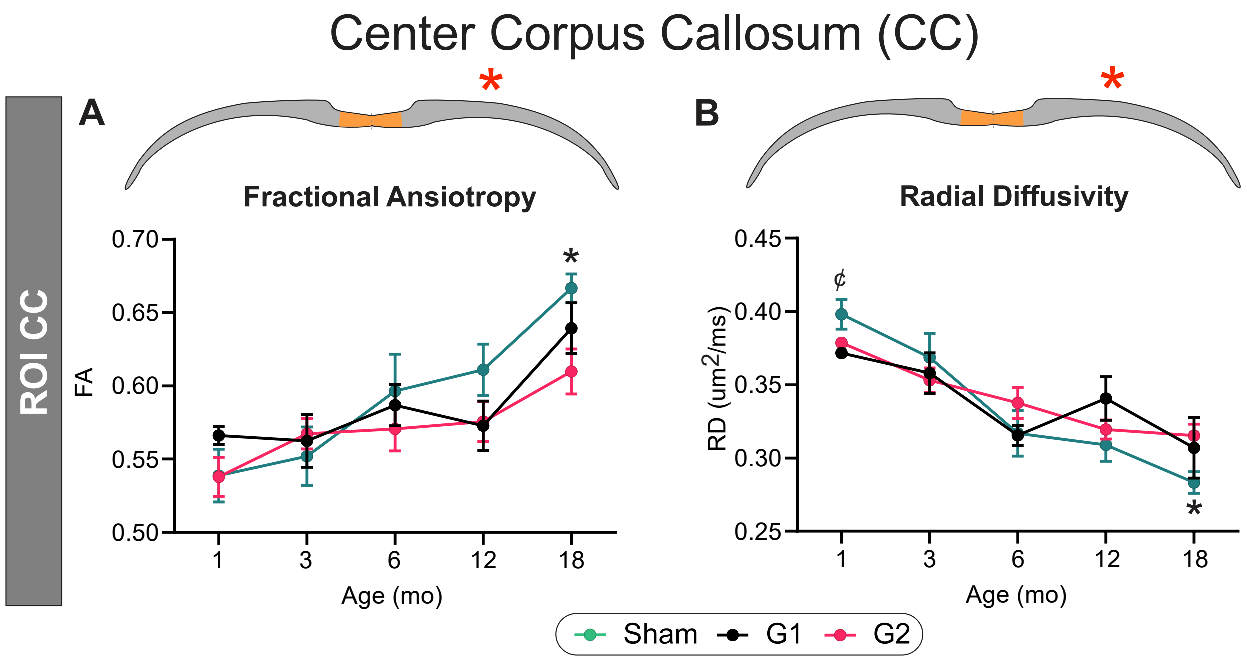


**Supplementary Figure 2. dMRI metrics from the midline (center) corpus callosum.** A) Fractional anisotropy of the center CC region of interest was significantly reduced in G2 mice at 18mpi (* $p<0.05$). B) Inversely, radial diffusivity (RD) was significantly increased in G2 mice at 18mpi (* $p<0.05$). G1 and Shams exhibited a trending difference at 1mpi (¢ $p<0.1$).

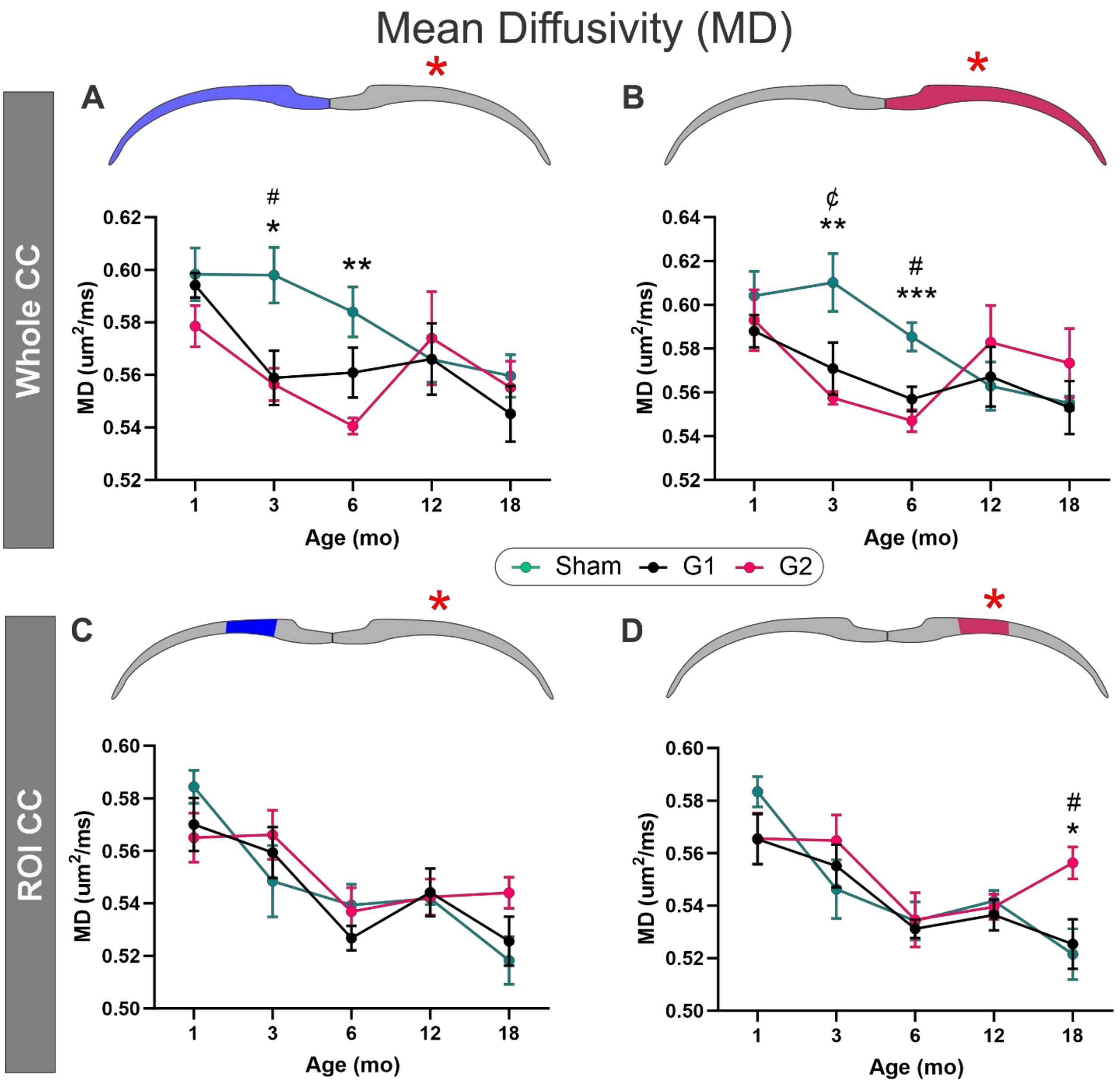


**Supplementary Figure 3. Mean diffusivity was altered in the corpus callosum in mid-life.** A) Whole CC mean diffusivity (MD) was significantly reduced in the contralateral CC with reductions at 3 and 6mpi in G2 mice compared to Shams. G1 mice were only significantly reduced at 3mpi. B) Identical to the contralateral CC there were significant differences within the whole CC with decreased MD at 3-6mpi compared to shams in both G1 and G2 mice. C) In regional CC measures no significant changes were found in the contralateral CC. D) There were also no significant ipsilateral MD changes in the regional CC measures except for G2 mice which that exhibited elevated MD at 18mpi, compared to G1 and sham mice. (* $p<0.05$, ** $p<0.01$, *** $p<0.001$ for G2 compared to shams; # $p<0.05$ for G1 compared to shams; ¢ $p<0.05$ for comparisons between G1 and G2)

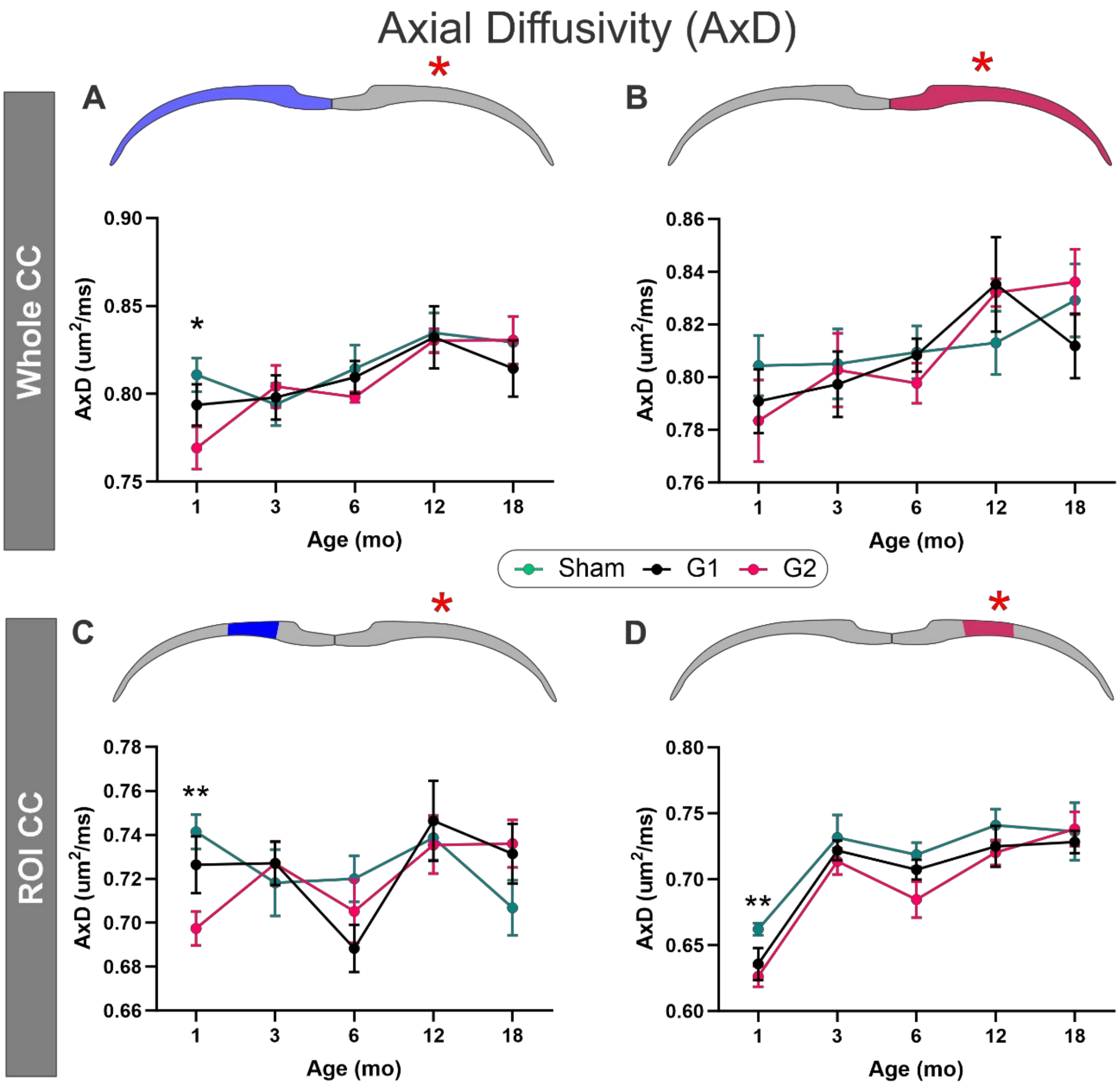


**Supplementary Figure 4. Axial diffusivity (AxD) was reduced in the corpus callosum at 1mpi.** A) Whole CC AxD was significantly reduced at 1mpi in G2 mice compared to Sham and G1 mice. B) No significant changes in AxD were observed on the ipsilateral CC at any timepoint, albeit G2 mice were reduced compared to shams (note increased variance at 1mpi). C) In regional CC measures only the G2 mice at 1mpi has significant reductions in the contralateral CC. D) Identical reductions in AxD were seen at 1mpi in the ipsilateral CC of G2 mice but not shams nor G1 mice. (* $p<0.05$, ** $p<0.01$ for G2 compared to sham)

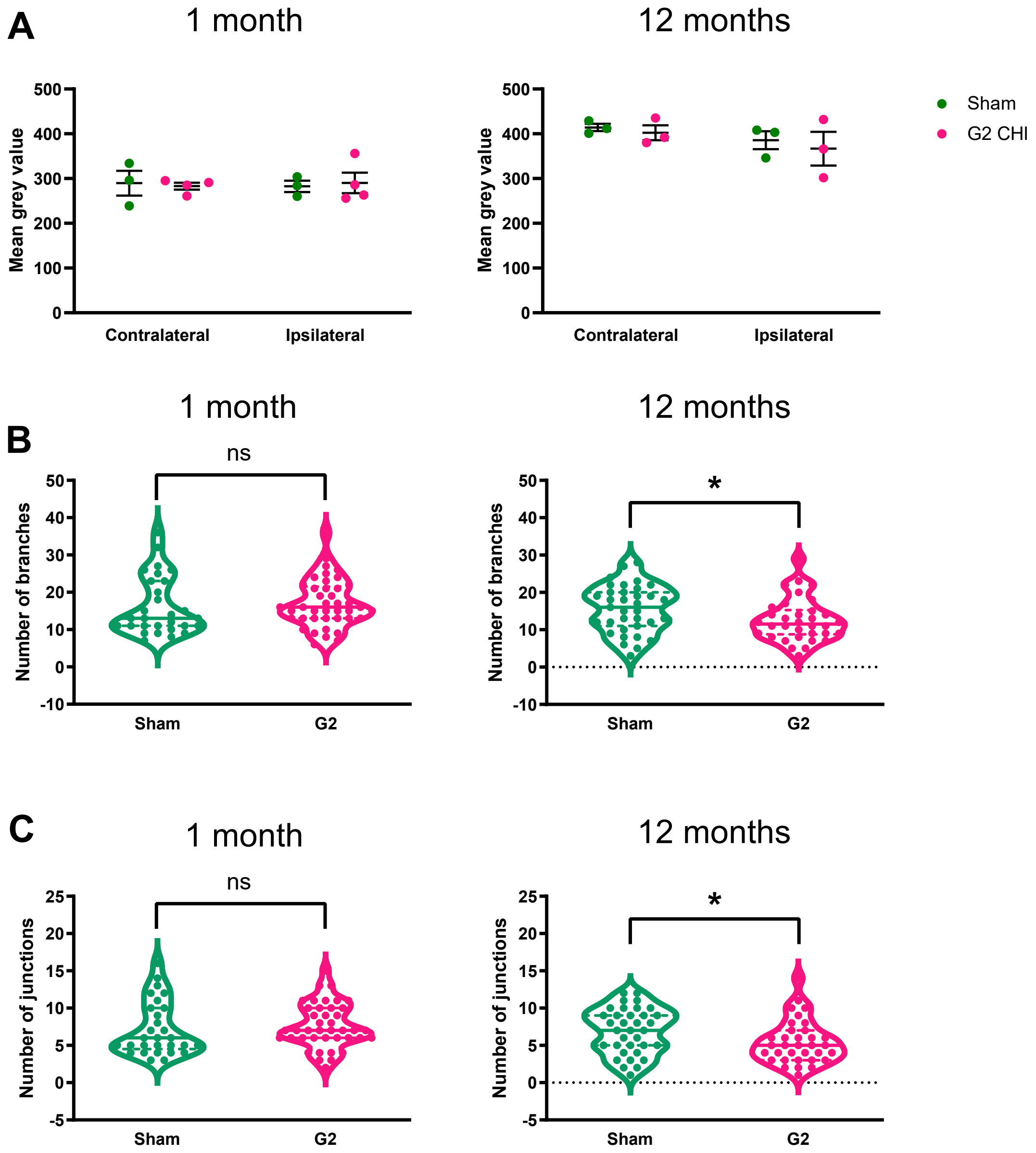


**Supplementary Figure 5. Long-term morphological changes in astrocytes after closed head injury (CHI) in the corpus callosum (CC).** Brain sections from sham and Grade 2 (G2) CHI mice were stained with anti-GFAP antibody. A) Mean grey value quantification showed no significant differences between sham and G2 groups at 1 and 12mpi (Kruskal-Wallis test, n=3-4 mice per group, data expressed as mean ± SEM). B) Astrocyte skeleton morphology analysis in lateral CC ROI showed a significant decrease of number of branches at 12mpi, but not at 1mpi (unpaired t-test, n=3-4 mice per group). C) Number of junctions in astrocytes was significantly reduced in G2 mice compared to sham group at 12mpi. There was no effect on the number of junctions at 1mpi (unpaired t-test, n=3-4 mice per group). * $p<0.05$

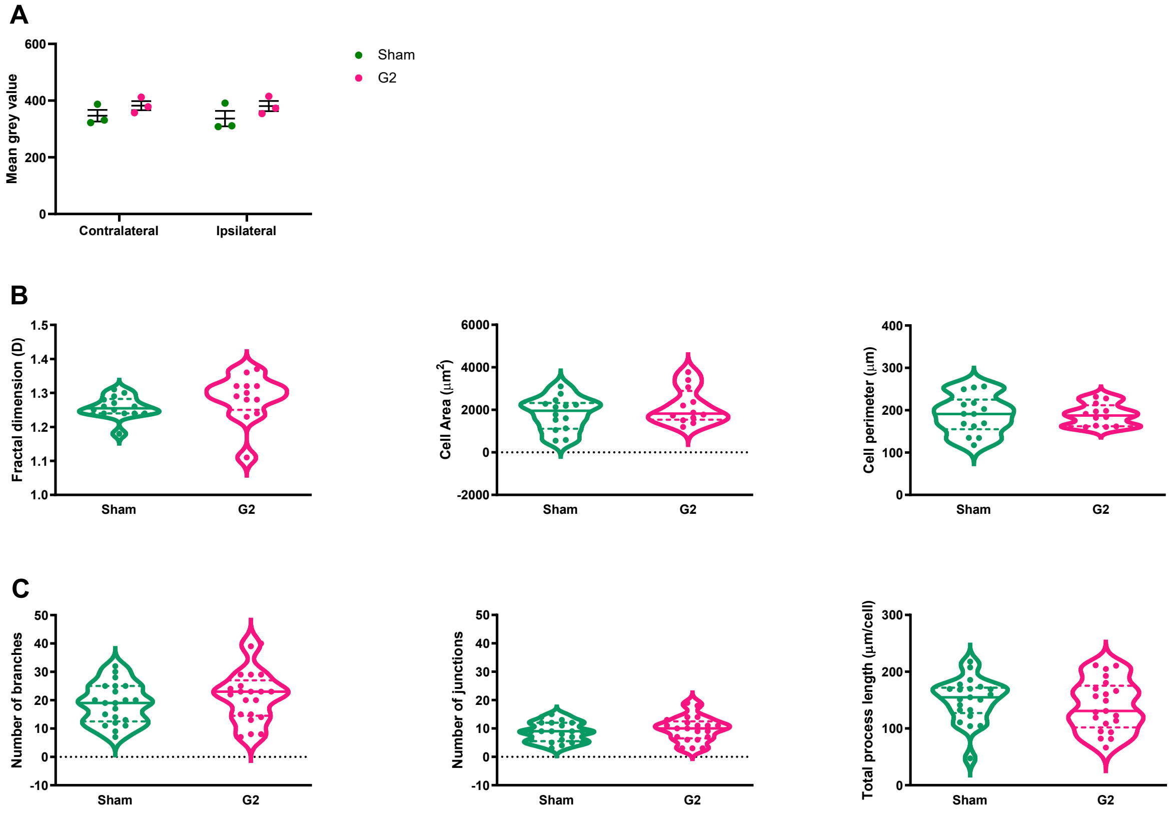


**Supplementary Figure 6. Effect of closed head injury (CHI) on microglial morphology in the corpus callosum (CC) at 12 months post impact.** A) Mean grey value quantification showed no significant difference between sham and G2 groups at 1 and 12mpi (Kruskal-Wallis test, n=3-4 mice per group, data expressed as mean ± SEM). B) Fractal analysis in ipsilateral CC found no significant differences between sham and G2 microglia in morphometric parameters such as fractal dimension (represents cell complexity), IBA1-positive cell area and cell perimeter (unpaired t-test, microglial cells pooled from 3-4 mice per group). C) Skeleton morphology analysis did not reveal significant differences in the number of branches, junctions and total process length (unpaired t-test, microglial cells pooled from 3-4 mice per group)

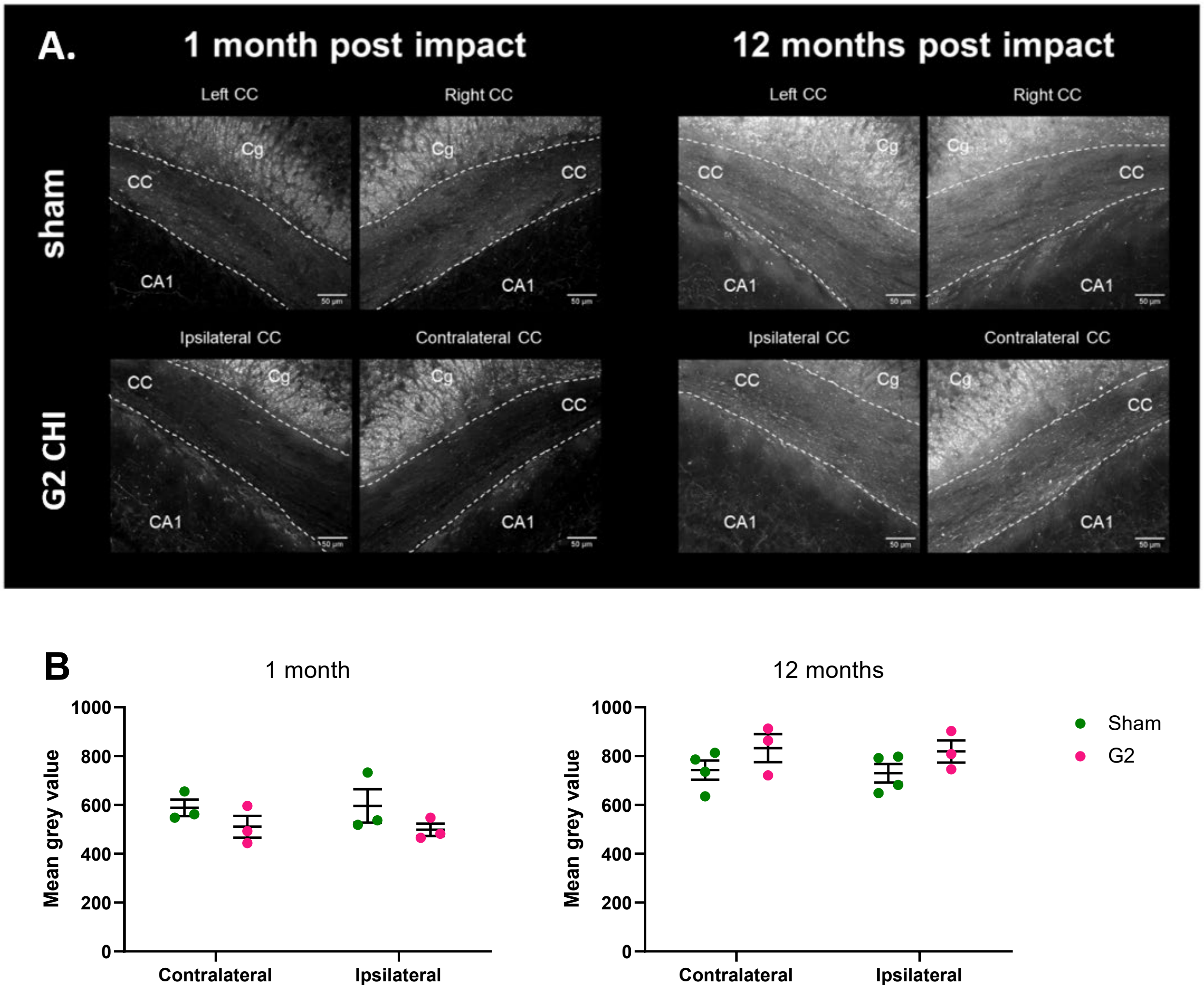


**Supplementary Figure 7. Neuronal staining quantification in the corpus callosum (CC) after closed head injury (CHI).** A) Coronal brain sections from sham (top) and Grade 2 (G2) CHI mice (bottom) stained with neurofilament 200 (NF200) marker for myelinated fiber neurons at 1 (left) and 12mpi (right). B) Mean grey value quantification showed no significant difference between sham and G2 groups at 1 and 12mpi (Kruskal-Wallis test, n=3-4 mice per group, data expressed as mean ± SEM).
